# Frontolimbic Functional Connectivity Development from Birth to Emerging Adulthood

**DOI:** 10.64898/2026.07.12.737639

**Authors:** Wonyoung Kim, Sarah Whittle, Andrew Zalesky, Ye Ella Tian

**Author notes:** Corresponding Authors: Wonyoung Kim, M.A. Department of Psychiatry University of Melbourne, 161 Barry St, Carlton, Victoria, 3053, Australia Ye Ella Tian, Ph.D.,Department of Psychiatry University of Melbourne, 161 Barry St, Carlton, Victoria, 3053, Australia.

## Abstract

Frontolimbic circuits connecting the frontal cortex with the amygdala and the hippocampus are critical for emotion and learning. These connections undergo major changes throughout childhood and adolescence, yet the principles guiding their maturation remain unclear. Analyzing large developmental cohorts, we show that fronto-hippocampus connectivity develops rapidly in early childhood and stabilizes thereafter. In contrast, fronto-amygdala connectivity becomes progressively differentiated from fronto-hippocampus connectivity, with development most pronounced during adolescence. This entailed hippocampus and amygdala respectively becoming preferentially tethered to association-related regions and sensorimotor-related regions. Greater adversity exposure was associated with more differentiated fronto-amygdala connectivity for a given age. Higher cognitive ability was associated with less differentiated fronto-hippocampus connectivity. These findings suggest that frontolimbic connectivity development follows a principle of differentiation, with circuit-specific timing and sensitivity to stress and learning. This work provides a foundation for understanding typical and atypical frontolimbic circuitry development from birth to emerging adulthood.

**Significance Statement:** The frontolimbic circuit, especially the fronto-hippocampus and fronto-amygdala circuitry, is integral for cognitive and affective processes. Yet its maturation has been challenging to characterize due to variable connectivity growth across the frontal cortex. Here, we reveal that this variability follows a principle of differentiation, where the hippocampus and the amygdala acquire respectively unique connectivity profiles with the frontal cortex at disparate developmental stages. Deviations from these normative trajectories were linked to cognitive ability and cumulative adversity, respectively. This approach provides a framework to unify inconsistent findings while serving as a foundation for identifying circuit-specific windows of plasticity and vulnerability.

## Introduction

The frontolimbic circuitry, specifically the frontal cortex functional connectivity with the amygdala and the hippocampus, supports a wide range of cognitive and affective functions. This includes approach/avoidance behavior, fear extinction, episodic memory, emotion regulation, social decision making, and stress regulation (1–6). Fronto-hippocampus and fronto-amygdala connections undergo substantial reorganization across childhood and adolescence (7–9) and have been implicated in a range of mental health problems (10–12). As such, understanding the development of the frontolimbic functional connectivity is critical for elucidating windows of plasticity and vulnerability (13–15).

Despite extensive research on fronto-hippocampus and fronto-amygdala circuitry development, a principle governing the spatial organization of frontolimbic connectivity development remains elusive. Prior studies on fronto-amygdala connectivity development report age-related increase from 4 to 23 years of age and from 7 to 22 (16, 17), decrease from 10 to 25 (18), and nonlinear decrease-then-increase from 7 to 24 (19) across the frontal cortex. Fronto-hippocampus connectivity growth patterns also vary by frontal cortex subregion with increase from 8 to 32 only in ventromedial prefrontal cortex but no change in ventrolateral or dorsolateral prefrontal cortex (7). Some report no age-related change in fronto-hippocampus connectivity from 4 to 10 and from 9 to 15 (20, 21). Importantly, recent studies show connectivity patterns in infancy to be distinct from those in adulthood for both fronto-hippocampus and fronto-amygdala connectivity (22, 23). When studied in isolation, however, such findings remain fragmented within the continuous developmental range from birth to adulthood. Together, these reports depict fronto-hippocampus and fronto-amygdala connectivity development as spatially variable across regions of interest and sample age ranges, calling for a coherent explanation across a continuous developmental trajectory.

Organizational principles have been shown to help synthesize disparate findings of connectivity development into interpretable knowledge. Organizational principles include global brain networks becoming more segregated (24, 25) or connectivity changing along the cortical hierarchy (26, 27). Such knowledge is also more interpretable when applied to human behavior or mental health. For instance, a landmark study demonstrated a principle where connectivity shows more dynamic developmental changes in association cortex regions than in primary cortex regions (28). Female-specific expression of this developmental principle was related to risk of depression (29), an insight that is more interpretable than identifying multiple connectivity measures associated with depression.

Connectivity differentiation is one such candidate principle. Differentiation is defined as connectivity profiles of a circuit becoming unique compared to another. Regions that begin with shared cortical connectivity patterns diverge in development to establish specialized cortical circuits (17, 30, 31). This principle is highly relevant to the hippocampus and the amygdala because they are linked to disparate functional brain networks: the Default Mode Network (DMN) and the Salience Network (SN), respectively (32–34). These networks each support externally directed and internally directed processes (35, 36). The two networks become increasingly segregated during normative development, a process thought to be critical for cognition and stress regulation (37–39). It is therefore possible that fronto-hippocampus and fronto-amygdala connectivity follows a similar principle of differentiation.

Individual deviations from typical fronto-hippocampus and fronto-amygdala maturational trajectories may hold implications for function and behavior. It has been suggested that the experience of adversity leads to advanced maturation of frontolimbic connectivity to help cope with the environmental demands (4, 40–42). While such an adaptation may be beneficial in the short term, it has been linked to long-term risk of psychopathology (10, 43). Cognitive maturation, including development of problem solving, decision making, and episodic memory, has also been suggested to be associated with frontolimbic connectivity development (44–47). Adversity and cognitive ability are thus two key factors that may explain why individuals diverge from the normative developmental trajectory for fronto-hippocampus and fronto-amygdala circuitry.

Here, we use resting-state functional magnetic resonance imaging (fMRI) to investigate the development and differentiation of fronto-hippocampus and fronto-amygdala connectivity from birth to emerging adulthood in an integrated framework. In this work, we operationalize differentiation as the degree to which a circuit’s connectivity pattern more closely resembles its own adult reference pattern while diverging from the other circuit’s reference pattern. We test whether the fronto-hippocampus and fronto-amygdala circuits show progressively more distinct connectivity patterns with age (i.e., differentiation) and whether individual deviations from typical developmental trajectories in these circuits relate to adversity and cognitive ability (Fig. 1). Over development, fronto-hippocampus and fronto-amygdala circuits were hypothesized to differentiate, becoming more preferentially connected to the DMN and SN, respectively. We also hypothesized that cognitive ability and adversity exposure would relate to differentiation of fronto-hippocampus and fronto-amygdala connectivity. Through such a framework, we sought to offer insights into when and how the fronto-hippocampus and fronto-amygdala connectivity profiles are organized.

**Figure 1.**
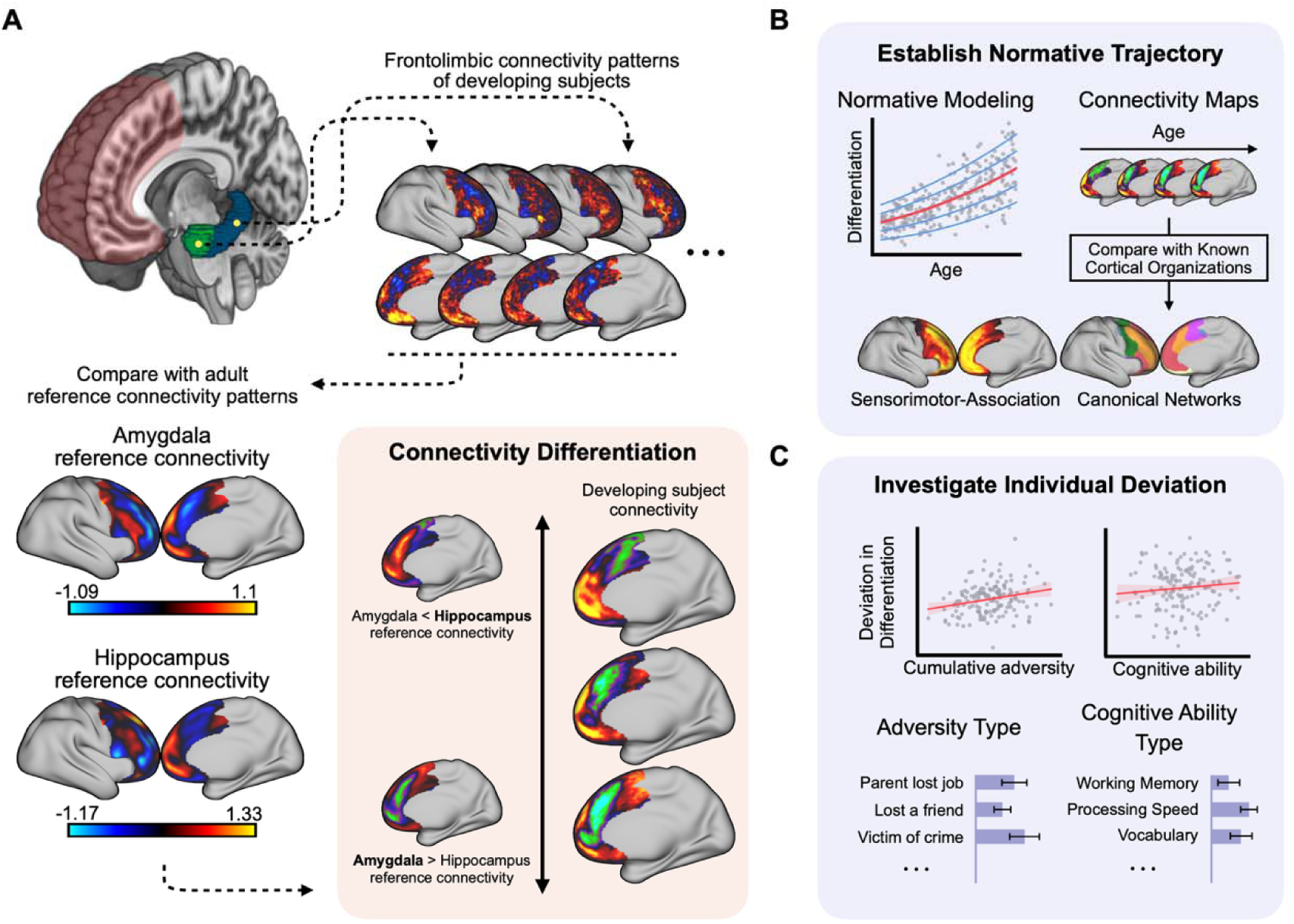
A conceptual overview of the study. A Connectivity of amygdala (green) or hippocampus voxels (teal) were calculated for across the frontal cortex (shaded red), which were then compared against adult reference connectivity patterns for the amygdala and the hippocampus. These connectivity maps followed a continuum of connectivity differentiation with amygdala-specific connectivity pattern on one end (bottom) and hippocampus-specific connectivity pattern on the other (top). B The normative developmental trajectory of connectivity differentiation was then mapped for fronto-amygdala and fronto-hippocampus connectivity. Normative modeling established the maturational trend over age. Connectivity map changes across age were also characterized and compared with known cortical organization schemes. C Individual deviations from the normative trajectory were inspected for functional implications including adversity and cognitive ability. Importance of adversity type and cognitive ability type was further probed.

## Results

We mapped the development of fronto-hippocampus and fronto-amygdala connectivity from birth to emerging adulthood in a large cross-sectional development cohort (N=822, age range 0-21 years) sourced from the Baby Connectome Project (BCP, N=195, 0-5 years) and the Human Connectome Project – Development (HCP-D, N=627, 5-21 years). Briefly, we first calculated the spatial correlation between the frontal connectivity patterns of a young person’s hippocampus or amygdala voxel and a group-average reference fronto-hippocampus and fronto-amygdala connectivity pattern estimated in healthy young adults (N=1,084, ages 22-35) (*SI Appendix*, Table S1). Then, we derived the differentiation index, the difference between the resemblance toward the reference hippocampus pattern and the reference amygdala pattern for each voxel (see Materials and Methods). Using this differentiation index, we characterize normative trajectories of fronto-hippocampus and fronto-amygdala differentiation and how they are represented in the hippocampus, the amygdala, and the frontal cortex.

### Normative Trajectory of Differentiation in Fronto-hippocampus and Fronto-amygdala Connectivity

First, we sought to model the developmental trajectory of the differentiation index to establish the normative development for both the hippocampus and amygdala. Fronto-hippocampus and fronto-amygdala connectivity became differentiated from birth to emerging adulthood at dissociable timings (Fig. 2A). Specifically, we found that fronto-hippocampus connectivity showed marked differentiation during early childhood (β = 0.239, p < 0.001), whereas fronto-amygdala connectivity differentiation was more prominent from late childhood to early adulthood (β = 0.371, p < 0.001). Direct comparison showed fronto-hippocampus differentiation to achieve 75.8% of its developmental range by the age of 1 year, whereas fronto-amygdala reached 4.2% at the same point (Fig. 2B, *SI Appendix, Methods*). Model specification and model fitting diagnostics are available in *SI Appendix*, Fig. S1 and *SI Appendix*, Table S2.

**Figure 2.**
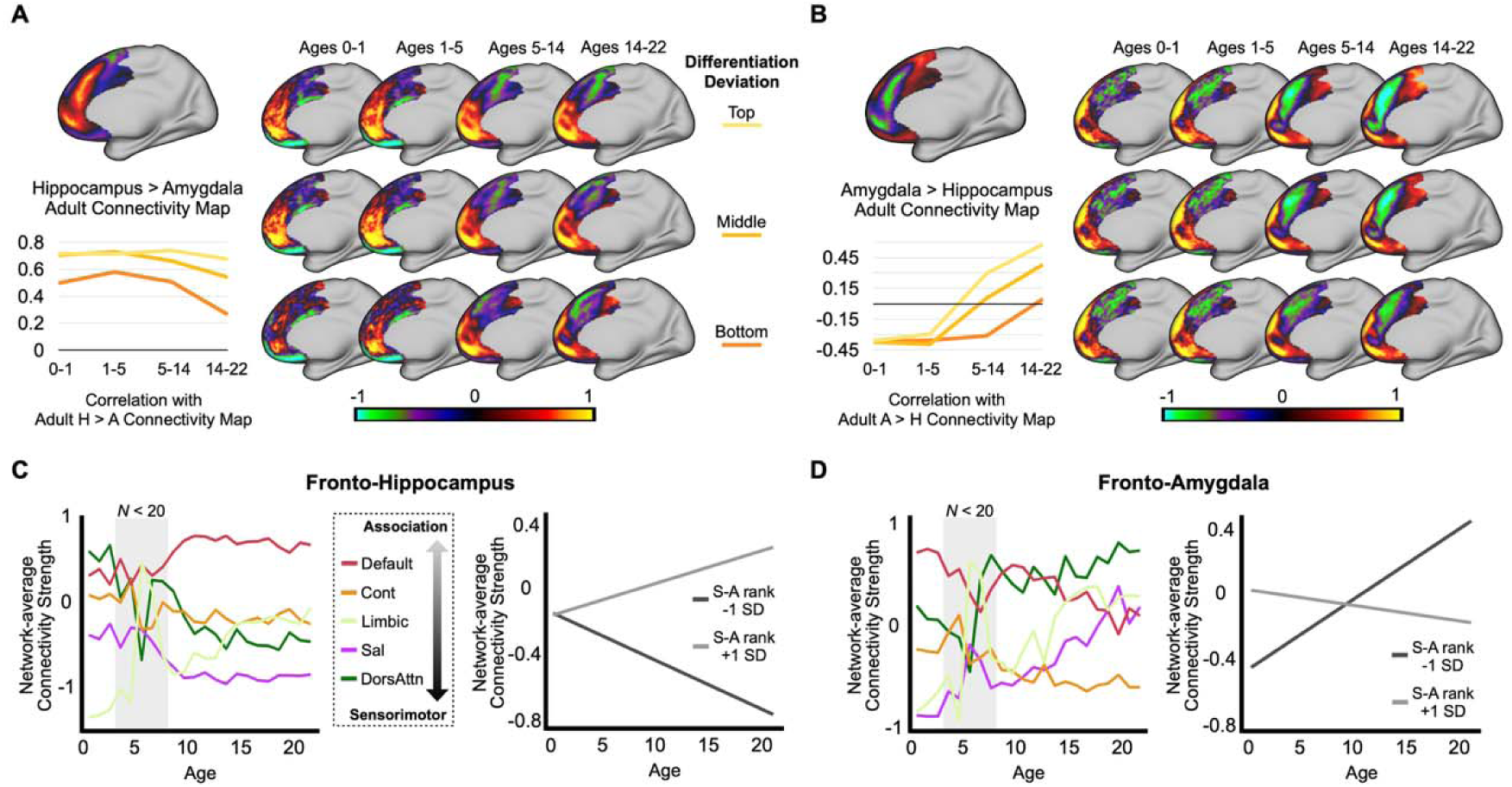
Differentiation of fronto-amygdala connectivity and fronto-hippocampus connectivity from birth to emerging adulthood. **A** The differentiation index was averaged within the hippocampus and within the amygdala and then fitted with GAMLSS modeling to characterize the normative trajectory. Red line shows the median while blue lines show centiles at 5%, 25%, 75%, and 95% from the top. From birth to emerging adulthood, amygdala and hippocampus connectivity became differentiated at different developmental timings. Piecewise linear effect testing reports the standardized slopes and significance for distinct developmental timings. **B** Each curve depicts the cumulative developmental progress made for each circuit’s differentiation. Fronto-hippocampus differentiation, drawn in solid cyan, shows pronounced early development followed by a slowed growth. Fronto-amygdala differentiation, drawn in solid vermillion, demonstrates gradual development that is protracted into adulthood. As an illustration, the percentage of development finished at age of 1 year is marked with dashed lines for both fronto-hippocampus and fronto-amygdala differentiation. **C** Voxels in the hippocampus (Hipp) show increasing connectivity differentiation in early childhood whereas the age-related increase in differentiation is pronounced in the amygdala (Amyg) during late childhood to emerging adulthood.

Next, we examined where differentiation occurs within the hippocampus and amygdala, given previously reported evidence of distinct roles of subregions in both structures (17, 20, 31). We first found six clusters dispersed across the left and right hippocampus during early childhood and three clusters of voxels in bilateral amygdala during late childhood to emerging adulthood (p_FWE-corrected_ < 0.05), confirming age-specific effects in the hippocampus and amygdala (Fig. 2C). Spatial permutation tests then revealed that age-related differentiation was not circumscribed to any established subregion of the hippocampus or the amygdala. This indicates that connectivity differentiation occurred at the level of the hippocampus and the amygdala, not specific subregions (see Materials and Methods, *SI Appendix*, Tables S3-6).

However, further exploration of continuous differences in age-related differentiation rather than within discrete subregion boundaries revealed that both the hippocampus and amygdala significantly differ along spatial axes in their developmental differentiation (*SI Appendix,* Fig. S2). In detail, the more posterior and medial the hippocampus voxels were, the more strongly they became differentiated toward the adult hippocampus reference over age (*SI Appendix,* Table S7). Simultaneously, the more anterior and lateral the amygdala voxels were, the more strongly they became differentiated toward the adult amygdala reference over age. This signifies that fronto-hippocampus and fronto-amygdala connectivity differentiation over development is not homogeneous but varies continuously within each structure.

Next, we asked how frontal cortex connectivity evolves across maturation to drive frontolimbic differentiation. Fig. 3A-B shows hippocampus and amygdala connectivity patterns to medial frontal cortex for multiple age bands (also shown in *SI Appendix*, Movies S1-2). For hippocampus connectivity, negative connectivity at dorsal-posterior regions and positive connectivity at ventral-anterior regions of the medial frontal cortex emerges early (Fig. 3A). These patterns show stability throughout childhood and adolescence. For amygdala connectivity, the same negative connectivity at dorsal-posterior regions moves toward ventral-anterior regions (Fig. 3B). This suggests development toward a configuration of connectivity differentiated from the hippocampus. These configurations respectively approached the adult reference hippocampus-specific or amygdala-specific connectivity maps (i.e., the difference between the amygdala and the hippocampus reference connectivity maps) throughout development. Connectivity across the lateral frontal cortex also showed age-related changes. Connectivity shifts in the lateral frontal cortex and across one-year age-bands are shown in *SI Appendix*, Figures S3-7 and *SI Appendix*, Movies S3-4.

**Figure 3.**
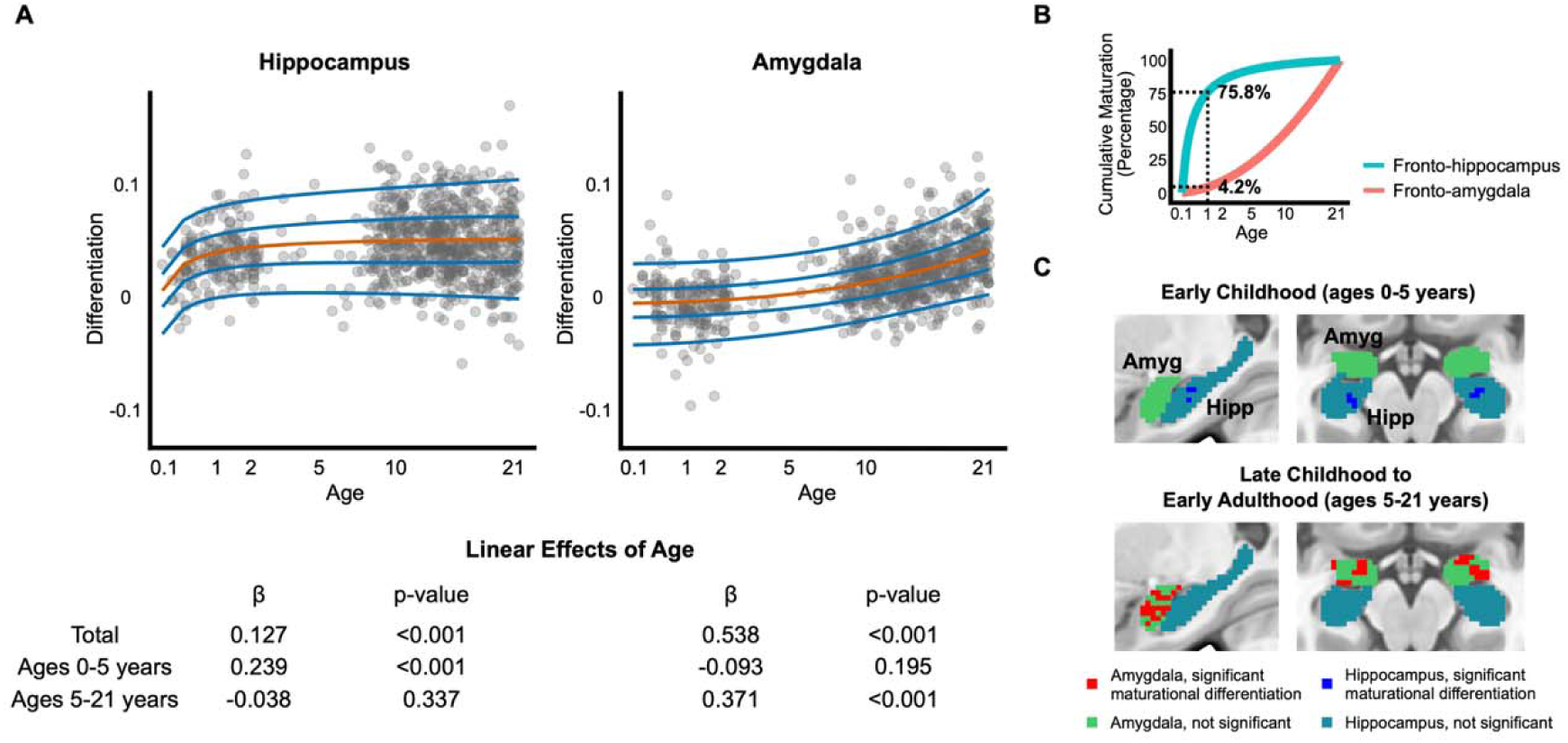
Characterization of differentiation in frontolimbic connectivity patterns. **A** Fronto-hippocampus connectivity is depicted, where the entire sample was divided into four age groups and within each age groups top, middle, and bottom tertile groups based on the deviation in hippocampus differentiation. Hippocampus > Amygdala adult connectivity map shows in the frontal cortex where hippocampus connectivity is stronger than the amygdala connectivity. The bottom left chart quantifies how similar the group average fronto-hippocampus connectivity pattern is to the adult Hippocampus > Amygdala connectivity map as a function of age and hippocampus differentiation deviation. **B** Same as panel A, fronto-amygdala connectivity is shown stratified by age and amygdala differentiation deviation. **C** Left panel depicts fronto-hippocampus connectivity averaged within each canonical network across the frontal cortex to show the age trend in network connectivity strength. Each point across age represents one-year age band group average connectivity maps. The canonical network labels are ordered from top to bottom in their relative position in S-A axis hierarchy. Area shaded in grey indicates potentially unstable estimations due to low subject count (*N* < 20 within a one-year age band). Right panel shows networks lower in S-A rank (-1 standard deviation, darker grey) decrease in connectivity with the hippocampus across age whereas networks higher in S-A rank (+1 standard deviation, lighter grey) increase. **D** Same as panel C, fronto-amygdala connectivity across frontal cortex networks and S-A axis hierarchy are mapped across age. Networks lower in S-A hierarchy increase in connectivity with amygdala while networks higher in S-A rank decrease.

To further understand the neurobiological implications of the connectivity shifts in the frontal cortex, we investigated the fronto-hippocampus and fronto-amygdala group-level connectivity maps in relation to known cortical organizations. We utilized the 7 canonical functional networks (48) and the Sensorimotor-Association (S-A) cortical hierarchy (49) that each mark discrete and continuous functional differences for cortical regions. Over maturation, fronto- hippocampus connectivity progressively aligned with association-related network regions (Fig. 3C, β = 0.156, p < 0.001) while fronto-amygdala connectivity aligned with sensorimotor-related network regions (Fig. 3D, β = -0.165, p < 0.001). Here, we tested if a network’s position within the S-A hierarchy (ranging from association-related to sensorimotor-related) was related to the slope of connectivity change (see Materials and Methods). Crucially, the results held even when leaving out the SN, the DMN, or both (*SI Appendix*, Fig. S8). This suggests that fronto-hippocampus and fronto-amygdala differentiation is bound to the functional hierarchy of frontal cortex regions.

### Adversity and Cognitive Ability are Linked to Individual Deviations in Fronto-hippocampus and Fronto-amygdala Differentiation

We next sought to understand whether deviation from the normative trajectory is associated with adversity and cognition. The extent to which an individual’s maturation in fronto-amygdala differentiation deviates relative to same-age peers was estimated (see Methods). Greater differentiation in connectivity than the norm at a given age was signified by a positive z-score, suggesting earlier maturation. Analyses were carried out in the late childhood to emerging adulthood cohort (i.e., HCP-D) where adversity and cognitive ability measures were available.

We found that greater deviation (i.e., higher z-scores) in the fronto-amygdala connectivity differentiation was significantly associated with greater cumulative exposure to adversity in the past year (r = 0.122, p = 0.003) (Fig. 4A). This suggests that adversity may be linked with more differentiated fronto-amygdala connectivity relative to same-aged peers. The association was not moderated by sex (p = 0. 353), age (p = 0.586), or socioeconomic status (p = 0.770), indicating that the effect of adversity exposure was consistent across sex, age, and socioeconomic status. Cumulative adversity was also associated with deviation in fronto-amygdala differentiation after controlling for socioeconomic status (p = 0.002), perinatal stress exposure (p = 0.026), and the state of stress at measurement (p = 0.002). This demonstrates that the adversity effect is not driven by how such factors may influence fronto-amygdala connectivity development (50–53). Deviation in hippocampus differentiation was not significantly associated with adversity exposure (r = 0.017, p = 0.671, Fig. 4A), which implies a circuit-specific role of adversity exposure. Next, we sought to understand the importance of adversity types as it is possible that a certain type of experience (e.g., related to threat or deprivation) may be driving the effect (54). To this end, principal component regression (PCR) was used to quantify the importance of 25 adverse life events in predicting the deviation in amygdala differentiation (see Materials and Methods and *SI Appendix*). No item was significant after multiple comparison correction, implying that fronto-amygdala differentiation may not be sensitive to specific adversity types beyond cumulative exposure to stress (Fig. 4B, *SI Appendix* Table S8). Adversity type importance was not attributable to the rate of endorsement (r = 0.109, p = 0.604), suggesting importance regardless of how many subjects in the sample had experienced them.

**Figure 4.**
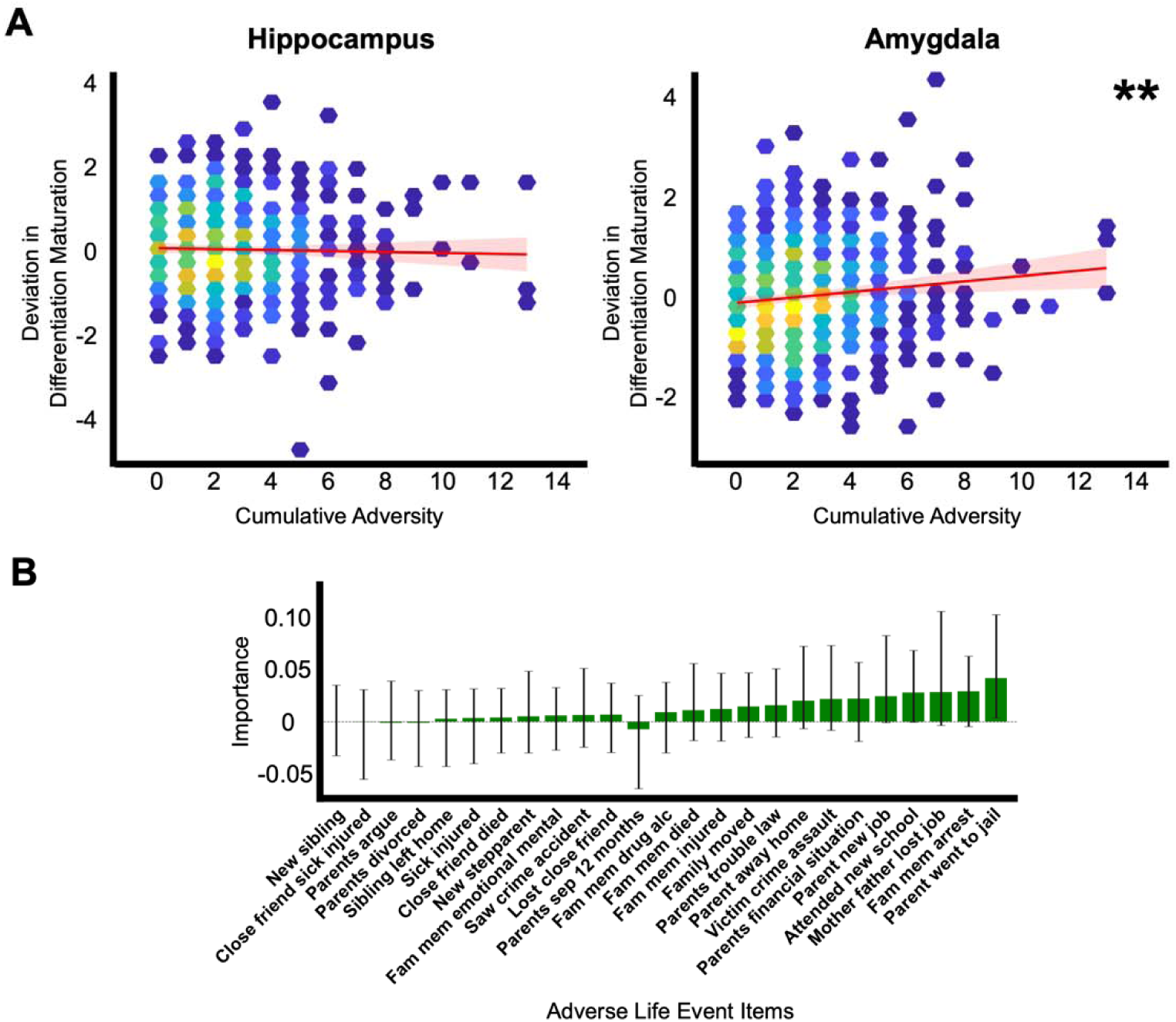
Link between adversity and individual deviation in fronto-amygdala differentiation. **A** Amygdala differentiation deviation, but not hippocampus differentiation deviation, was positively correlated with cumulative adversity. Colors of data points indicate density, where brighter and warmer colors show higher count of individual overlap. Red line shows linear fit between the two variables with 95% confidence interval shaded in brighter red. **B** In principal component regression models that predict with the 25 adverse life event items the deviation in amygdala differentiation maturation, importance was calculated for each item. Whiskers overlaid on each bar show 95% confidence intervals, where not including zero suggests significance before multiple comparison correction. Significance indicated as **p < 0.01.

We then investigated the association between cognitive ability and the maturation of frontolimbic differentiation. We found that the total age-adjusted cognitive ability score was associated with lesser fronto-hippocampus differentiation (Fig. 5A, r = -0.138, p = 0.003). Cognitive ability score summarized capacity in episodic memory, cognitive flexibility, inhibition, language decoding, vocabulary comprehension, processing speed, and working memory. This suggested that less differentiated fronto-hippocampus connectivity was associated with higher cognitive ability at a given age. Lack of interaction with sex (p = 0.567), age (p = 0.609), and socioeconomic status (p = 0.163) suggested a relationship between fronto-hippocampus differentiation and cognitive abilities general to sex, age, and socioeconomic status. Results remained after controlling for socioeconomic status, the state of stress at measurement, or perinatal stress exposure (p = 0.008; p = 0.005; p =0.007), showing that the effect is not confounded by these factors (50, 51, 55). Deviation in amygdala differentiation development was not correlated with cognitive ability (r = 0.033, p = 0.487).

**Figure 5.**
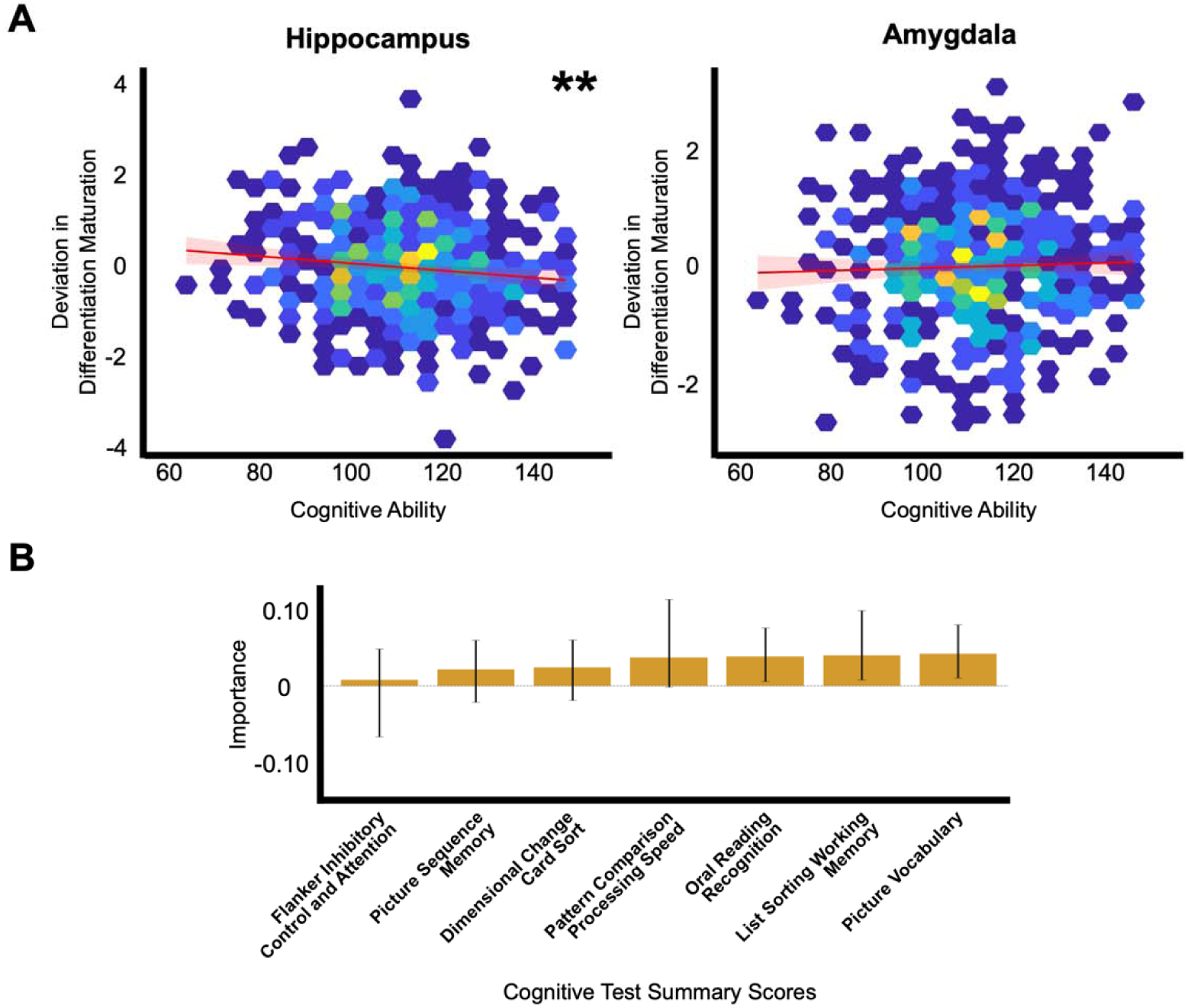
Link between cognitive ability and deviation in fronto-hippocampus differentiation maturation. **A** Hippocampus differentiation deviation, but not amygdala differentiation deviation, was negatively correlated with age-adjusted cognitive ability summary score. **B** In principal component regression models that predict the deviation in hippocampus differentiation with the 7 cognitive test scores, predictive importance was calculated for each test score. Whiskers overlaid on each bar show 95% confidence intervals, where not including zero suggests significance before multiple comparison correction. Significance indicated as **p < 0.01.

Next, we investigated the importance of cognitive domains that were important in fronto-hippocampus differentiation deviation as it is possible that certain domains of intelligence (e.g., episodic or working memory) may be potentially driving the effect (47). The same PCR procedure carried out in the adversity type importance analyses was employed, revealing importance of oral reading recognition, list sorting working memory, and picture vocabulary after multiple comparison correction (Fig. 5B, *SI Appendix* Table S9). All cognitive scores were age-adjusted. Among these, picture vocabulary and oral reading recognition are considered to gauge intelligence learned through life experiences whereas other tests measure innate intelligence (56). Such results therefore indicated that lesser fronto-hippocampus differentiation was relatively more related to cognitive ability and skills accrued through life experiences rather than innate abilities such as episodic memory or processing speed.

### Replication Analyses

To assess the generalizability of our findings, we sought to replicate the core findings in a community sample of developing individuals (Philadelphia Neurodevelopmental Cohort, PNC, N = 898, ages 8-23). We showed that the pattern of fronto-hippocampus and fronto-amygdala connectivity differentiation across development was replicated (r = 0.266, P_brainSMASH_ < 0.001). In detail, connectivity of amygdala voxels increasingly resembled adult fronto-amygdala connectivity more than fronto-hippocampus connectivity and hippocampus voxel connectivity showed the opposite trend (*SI Appendix*, Fig. S9). However, the effect size was weaker than in the main analyses (*SI Appendix*, Fig. S10-11). One possibility is that the developmental effect on differentiation is weaker at the population scale than our main sample. Alternatively, higher individual variability and scan duration may have contributed to the modest effect sizes. Individuals with a variety of clinical and subclinical psychiatric symptoms were included in the PNC dataset (57), potentially introducing noise to the relationship between age and differentiation (10–12). Moreover, prior evidence suggests that individual-specific signal during resting-state becomes more reliable as scan duration increases from ∼5 minutes (i.e., scan length of the PNC dataset) to ∼25 minutes (i.e., scan length of the HCP-D dataset) (58, 59). Future studies on how differentiation measurement is impacted by disease effects and scan parameter effects may disambiguate such possibilities. Individual difference in fronto-amygdala differentiation in the PNC was correlated with lifetime exposure to traumatic events (r = 0.071, p = 0.034), providing converging evidence of fronto-amygdala differentiation implicated with adversity (*SI Appendix*, Fig. S12 and Table S10). Individual difference in fronto-hippocampus differentiation was not associated with cognitive ability (r = 0.015, p = 0.664), potentially owing to limited representation of learned intelligence in the cognitive measures of the PNC or lower generalizability of the cognitive ability association relative to the adversity association.

### Sensitivity Analyses

Lastly, we conducted multiple sensitivity analyses to test the robustness of our core findings. In brief, our results converged across different methods for computing the core differentiation metric and tests of specificity in key developmental and functional associations as well as scanning conditions. Details are reported in *SI Appendix*, Fig. S13-19.

## Discussion

In this work, we provide evidence that fronto-hippocampus and fronto-amygdala connectivity follows the principle of differentiation from birth to emerging adulthood. The circuits examined here are anatomically more accurately described as mesiotemporal–prefrontal connections. However, for consistency with existing literature, we use the broader term “frontolimbic connectivity,” by which we specifically mean fronto-hippocampus and fronto-amygdala circuits. At birth, the hippocampus and amygdala share a common connectivity blueprint with the frontal cortex. Fronto-hippocampus connectivity becomes unique from the amygdala’s from birth to early childhood, with fronto-amygdala connectivity diverging from hippocampal patterns later in development. Such circuit-specific timings of development indicate that frontolimbic system growth entails not only maturation of respective structures but also system-level reorganization along a principle of differentiation. This is also likely related to how different cognitive and affective functions emerge at different timings. Our findings converge to support this idea as well, where frontolimbic circuitry differentiated towards the opposite ends of the sensorimotor-association axis while showing mutually exclusive associations with adversity and cognition. Connectivity differentiation provides an explanation for how frontolimbic connectivity development is spatially organized.

Our results demonstrate that differentiation occurs within the frontolimbic system across development. Our findings of within-system differentiation might initially appear to contradict the notion of the fronto-hippocampus and fronto-amygdala circuitry functioning as an integrated system. Rodent studies of fear extinction show that hippocampus and amygdala are sequentially recruited across development to form a functional system for fear extinction (60). In humans, the hippocampus and the amygdala share a great amount of overlap in connectivity (9, 34, 61, 62), potentially supporting integrated functions such as emotion regulation or stress regulation (2, 4). Nonetheless, specialized connectivity patterns within the frontolimbic circuitry may indicate better integration of functionalities from diverse frontal cortex regions (63). Differentiation within the integrated fronto-hippocampus and fronto-amygdala system may thus be what enables complex behaviors to arise in development.

Importantly, differentiation unfolds along distinct timelines for fronto-hippocampus and fronto-amygdala connectivity. This may indicate dissociable windows of plasticity where life experiences could affect connectivity organization. This extends the existing models of frontolimbic development in volume and glucocorticoid stress response (4, 13, 43, 64). In these works, the hippocampus is fully organized by adolescence and is most malleable during the early postnatal years. In contrast, the amygdala shows protracted development and thus remains malleable into adulthood. Our results show that fronto-hippocampus and fronto-amygdala connectivity organization follows a similar timeline. Differential growth spurts of connectivity patterns may represent connectivity becoming specialized to support functions necessary at different stages of maturation (65). It may be such a functional specialization that makes the circuitries easily influenced by distinct types of life experiences (66).

Supporting the concept of functional specialization, our finding of frontolimbic connectivity differentiation was aligned with the cortical hierarchy across the frontal cortex. There is converging evidence that brain connectivity maturation is organized in a smooth gradient from sensorimotor regions at one end and association regions at the other (27, 28, 49, 67). Our results expand this framework, demonstrating that the previous reports of fronto-hippocampus and fronto-amygdala connectivity development may follow a differentiation occurring along the S-A axis. This may imply that the fronto-amygdala circuit becomes relatively more involved in attention coordination while the fronto-hippocampus circuit becomes more specialized for associative reasoning. Such a phenomenon nonetheless extends beyond the hippocampus-DMN and amygdala-SN connectivity dichotomy, involving frontal regions outside the two networks. It is therefore likely that fronto-hippocampus and fronto-amygdala connectivity, distributed across diverse hierarchical positions from sensorimotor-related to association-related, together support essential functions.

Furthermore, exploratory analyses showed that there are continuous gradients of developmental differentiation within the hippocampus and amygdala, respectively. In prior studies, distinct subregion-specific connectivity patterns emerged across development when divided into anterior/posterior hippocampus and centromedial/basolateral amygdala (17, 20, 22). Our findings extend this and show developmental connectivity differentiation occurring both across and within the two structures. This unifies previous accounts of within-structure connectivity gradients with our fronto-hippocampus and fronto-amygdala differentiation framework (9, 68, 69). The direction of connectivity recapitulated the well-documented anterior-posterior axis of hippocampus aligning with the S-A axis while also revealing that the lateral-medial axis of amygdala follows a similar organization. This suggests that frontolimbic differentiation may also encompass developmental subregion- or gradient-level connectivity refinement that is purported to be involved in the formation of complex cognitive and affective functions (20, 68).

Further bolstering the functional implications of our findings, adversity was linked to higher differentiation in fronto-amygdala connectivity at a given age. One possible explanation is that the experience of stress led to deviations from the normal trajectory, though the cross-sectional nature of our individual deviation measures limits mechanistic or causal interpretations. This is in line with previous literature observing stress-related advancement in fronto-amygdala connectivity (40, 41, 70). Of note, accumulated evidence suggests distinct dimensions of stress exerting different effects on neurodevelopment (54, 71). Our findings showed the influence of adversity types to be limited. This suggests that the process of frontolimbic connectivity pattern shifts is likely susceptible to accumulated stress rather than specific types of stress. In parallel, threat- or deprivation-related stress may affect connectivity strength of more localized functional circuits (72, 73). The lack of association between adversity and fronto-hippocampus differentiation may also imply that after connectivity organization is established in early childhood, stress may impact the fronto-hippocampus local connectivity strength changes but not widespread pattern shifts (10).

In contrast, we found that higher cognitive ability was related to negative deviations of fronto-hippocampus differentiation. This may indicate that active learning may prolong a less mature state that promotes plasticity and learning, mirroring insights from cortical structure studies (74, 75). Cognitive domains of significant and higher importance revealed from our analysis, including oral reading recognition and picture vocabulary, are learned in nature. This supports such an experience-dependent interpretation of our findings. It should be noted that list sorting working memory, a test of relatively innate intelligence, was also of high importance. This implies that the contrast between inherent versus acquired intelligence is not discrete but rather a trend. Additionally, previous findings suggest that deprivation of learning environment, not enrichment, is associated with delay in brain maturation (54, 71). One possible explanation is that the transitional state of the hippocampus becoming more preferentially connected to association-related regions in the frontal cortex supports learning, thus making less mature states beneficial. Conversely, measures such as cortical thickness could more directly reflect development of functional capacity where delayed maturation is maladaptive. Probing fronto-hippocampus connectivity differentiation during cognitive tasks may further disambiguate such a possibility. Lastly, we found a lack of association between cognitive ability and fronto-amygdala differentiation deviation. While the fronto-amygdala circuit is also implicated in intelligence and learning (44), such impact may be confined to changes in local connectivity strength rather than the process of fronto-amygdala connectivity differentiation.

Some methodological limitations warrant discussion. First, there was a significantly smaller number of participants in the age range of 3 to 8, which restricts inferences made within this age. Though we believe that the trajectories we estimated within this age range are plausible, it should be clarified that they can be less reliable in age ranges where data is sparse. As this age range is difficult for data collection due to excessive movement (76), data collection efforts within this age range will greatly aid in understanding the fine-grained maturational shifts occurring at this age especially when considering its maturational significance (4, 13, 43, 64). Second, the cross-sectional nature of our data limits interpretations. Age trajectories of frontolimbic differentiation derived from cross-sectional data may underestimate how individuals truly age over time (77). Interpretations linking differentiation deviation with the degree of maturation thus require caution. Nonetheless, the overall direction or the distinct timings of change we found may be more reliable as partly validated with cohort-separate analyses in *SI Appendix*. Critically, it is equally plausible for individual deviations to exist before experience of adversity or learning. Evidence exists for both experience-dependent fronto-hippocampus and fronto-amygdala connectivity changes (51, 78, 79) and individual differences predicting future intelligence or vulnerability to adversity (80, 81). Longitudinal studies leveraging our differentiation framework could adjudicate such possibilities. Third, partial replication in a community dataset may indicate that applying our framework may be challenging at the population level with high individual heterogeneity. While this is a limitation that also confounds traditional connectivity frameworks especially when involving regions susceptible to noise, ongoing efforts to mitigate such confounds may alleviate such concerns in future studies (82). Fourth, the adult connectivity patterns may not represent a homogeneous developmental endpoint. Endpoints for brain development may significantly differ across individuals given the disparate developmental trajectories (75, 83). It is also possible that the circuits continue to evolve during early adulthood (25). Nevertheless, our supplementary analyses showed that the adult reference connectivity patterns are largely consistent over randomly sampled individuals and over three consecutive age-bin groups in early adulthood (*SI Appendix*, Fig. S16 and Table S11). In other words, our use of the adult connectivity is not heavily impacted by potential heterogeneous developmental endpoints within the adult sample for the purpose of indexing developmental change. Fifth, the associations between the differentiation deviation and adversity and cognitive ability were modest. Nonetheless, they may still inform specific developmental mechanisms that link frontolimbic circuit maturation to adversity and cognition. In detail, our findings highlight fronto-amygdala and fronto-hippocampus circuitry respectively and specifically linked to adversity and cognition. Despite the modest effect sizes, this may indicate that fronto-hippocampus and fronto-amygdala connectivity are functionally dissociable in the context of developmental reorganization of connectivity. Further studies are required to assess the robustness and specificity of these behavioral links. Sixth, inference on the timing effect of exposure to adversity is limited. As fronto-amygdala development demonstrates a distinct timing, exposure to adversity may be especially critical in some developmental phases (84). The lack of age moderation partially addresses such a possibility and implies that the effects were similar between younger and older subjects. Nonetheless, our data utilizes an adversity measure that is confined to past-year occurrences, which limits any claims on exposure timing beyond the past year. While collecting data on past-year adversity is less prone to retrospective errors, future studies should also collect data on past adversity exposure along with timings of exposure to further delineate the effect of adversity.

In conclusion, we demonstrate that fronto-hippocampus and fronto-amygdala connectivity development follows a principle of differentiation at circuit-specific timings. Individual deviation in connectivity differentiation was associated with adversity and cognitive abilities. The pattern of connectivity differentiation was hierarchically organized, with amygdala differentiating toward sensorimotor-related regions of the frontal cortex and the hippocampus differentiating toward association-related regions. Together, our work shows differentiation from birth to emerging adulthood as a framework for understanding normal and abnormal frontolimbic connectivity development.

## Materials and Methods

*SI Appendix* details data acquisition and preprocessing as well as statistical analyses. In short, we utilized resting-state scans from multiple large-scale datasets collected at different sites: the Human Connectome Project Young-Adult (HCP-YA) dataset (N*=*1,084; ages 22-35) (85), the Human Connectome Project - Development (HCP-D) dataset (N=627, ages 5-21) (86), and the Baby Connectome Project (BCP) dataset (N=195, ages 0-5) (87). The HCP-D and HCP-YA datasets are distributed minimally preprocessed and normalized to a standardized surface mesh using multimodal surface matching (88). The BCP dataset was preprocessed using Nibabies v25.2.1. (89), the infant-equivalent of fMRIPrep. To account for the distinct morphology of the developing brain, NiBabies utilizes age-appropriate preprocessing components such as infant-specific templates and Infant FreeSurfer (90, 91). Further details on preprocessing can be found in *SI Appendix, Methods*. Approval was obtained from respective institutional review board bodies that contributed to the datasets. Written informed consent was obtained from each participant and from parents.

### Normative Development of Fronto-hippocampus and Fronto-amygdala Connectivity Differentiation

We first calculated the differentiation index. The differentiation index (DI) was calculated for each voxel in an individual as follows:

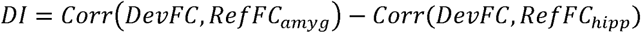

where a developing individual’s frontal cortex functional connectivity map of each amygdala or hippocampal voxel is denoted *DevFC (SI Appendix, Methods)*. *RefFC_amyg_* and *RefFC_hipp_*indicate the adult reference connectivity patterns, calculated as the HCP-YA group-average frontal cortex functional connectivity maps, averaged across all hippocampus or amygdala voxels. In detail, two spatial correlations were calculated per each voxel across all hippocampus or amygdala voxels: one measuring the similarity of their fronto-amygdala connectivity pattern to the reference adult fronto-amygdala connectivity pattern, and another for the hippocampus. The differentiation index was defined as the voxel-wise difference between these two correlation values. Because the differentiation index is directionally opposite for amygdala and hippocampus, differentiation indexes in the hippocampus were multiplied by -1 so that higher differentiation indexes in both the amygdala and the hippocampus track the degree of differentiation in the same direction (i.e., *DI = Corr(DevFC, RefFC_amyg_*) - *Corr(DevFC, RefFC_hipp_*) for amygdala voxels whereas *DI = Corr(DevFC, RefFC_hipp_*) - *Corr(DevFC, RefFC_amyg_*) for hippocampus voxels). Differentiation index was averaged for the hippocampus and the amygdala respectively for statistical analyses. An alternative definition of the differentiation index was also explored as described in *SI Appendix, Methods* and *Fig. S13*. Hippocampus and amygdala voxels were defined using the Melbourne Subcortex Atlas, while frontal cortex regions were extracted from the Glasser Atlas (92, 93)(*SI Appendix, Table S1)*.

To accurately model the normative trajectory of the average connectivity differentiation for the amygdala and the hippocampus, respectively, Generalized Additive Model for Location, Scale, and Shape (GAMLSS) models were fit (94). Intracranial volume and mean framewise displacement were regressed out of the amygdala and hippocampus average differentiation index prior to fitting. Age and sex were entered into the model with site also added as a random effect term. Information on model fitting is described in *SI Appendix, Methods*. Z-scores that quantify the individual deviations from the estimated norm was calculated in a 10-fold cross-validation scheme for each of the two normative models. The z-scores were used in all subsequent statistical analyses to characterize individual deviations. To ascertain that the hypothesized developmental circuit differentiation and the observed contrast in developmental timings are statistically significant, the effect of development was probed with linear regressions. In detail, we tested the linear effect of age for the entire sample as well as for subjects before and after the age of 5 years. Age 5 was selected as a descriptive breakpoint because it corresponded to the boundary between the two developmental cohorts, allowing us to test whether the observed shift in developmental timing was present within each cohort rather than being attributable to the transition between cohorts.

To fully characterize the spatial extent of the age effects on differentiation across the hippocampus and amygdala, we carried out discrete and continuous subregion analyses. We first examined the effect of age on the voxel-wise differentiation index maps of the individuals to identify voxels that became increasingly differentiated with age in frontal connectivity patterns, using *randomise* in FSL. Then, the amygdala was further parcellated into medial and lateral subregions, and the hippocampus was divided into anterior and posterior subregions (92). For the discrete subregion analysis, the t-statistic values of the age effects for a subregion were summed and compared. If a subregion had higher summary t-statistic value than the other, we treated this as evidence of discrete subregion-specific effect. For the continuous subregion analysis, we tested a multiple regression model with the t-statistic value as the dependent variable and coordinates in MNI space (mm) along the X axis (medial-lateral axis), Y axis (posterior-anterior axis), and Z axis (dorsal-ventral axis) as independent variables. If the t-statistic values change as functions of spatial axes, we treated this as evidence of continuous subregion-specific effect. Full details of the subregion analyses can be found in *SI Appendix, Methods*.

### Association with Adversity and Cognition

Pearson correlation was first used to test the association between the z-scores of fronto-hippocampus and fronto-amygdala differentiation and behavioral variables measuring adversity and cognition. Cumulative adversity was calculated as the sum of adverse life events experienced in the past year, as reported in a questionnaire as a part of the HCP-D data collection (95). Cognitive ability was measured through a composite of cognitive assessments in the NIH toolbox (96). Socioeconomic status, recent stress, or perinatal stress exposure were also included as covariates. To account for the association effects potentially varying across age, sex, or socioeconomic status, separate models were tested as these variables included as interaction terms.

To understand the contributing role of adversity items and cognitive domains, Principal Component Regression (PCR) models were utilized. PCR first identifies the strongest pattern of variation among a set of variables and then tests its association with the outcome variable. Bootstrapping was carried out 10,000 times to describe whether a feature was important within the PCR models, robust to potential sampling biases. Feature importance was calculated by aggregating the principal component loadings across the items that together predicted the differentiation z-score (mean r_predicted-actual_ = 0.133, 95% CI [0.02 0.23] for fronto-amygdala; mean r_predicted-actual_ = 0.135, 95% CI [0.02 0.25] for fronto-hippocampus). The significance for each item was assessed by examining the confidence intervals calculated through bootstrapping. If the 95% confidence interval of the principal component loading for an item did not contain 0, the item was deemed to be statistically significant, equivalent to a p-value being smaller than 0.05. Correction for multiple comparisons was performed using the False Discovery Rate (97).

## Data and Code

The HCP-YA dataset is publicly available through ConnectomeDB through an access request (https://db.humanconnectome.org). The BCP dataset and the HCP-D dataset can be accessed by submitting data access requests to the National Institute of Mental Health Data Archive (https://www.humanconnectome.org/study/lifespan-baby-connectome-project, https://www.humanconnectome.org/study/hcp-lifespan-development/data-releases). The PNC dataset is also available through a data access request (https://www.ncbi.nlm.nih.gov/projects/gap/cgi-bin/study.cgi?study_id=phs000607.v3.p2). Custom code for analyses and visualisation is shared at https://github.com/1youn9-kim/frontolimbic_fc_dev and archived on Zenodo (DOI: 10.5281/zenodo.18169371). Additional tools used to work with CIFTI files that were used to support the findings for this study were as follows: Connectome Workbench v1.5.0 (https://www.humanconnectome.org/software/connectome-workbench), R 4.4.0, and MATLAB 2023b-2024b (MathWorks, https://www.mathworks.com/). Processing neuroimaging data involved the use of fMRIPrep v25.1.1. (https://fmriprep.org/), Nibabies v25.2.1. (https://nibabies.readthedocs.io/en/latest/), and FSL 5.0 (https://fsl.fmrib.ox.ac.uk/fsl/fslwiki).

## Acknowledgments

W.K. is supported by a Melbourne Research Scholarship and a Nick Christopher Scholarship at the University of Melbourne. Y.E.T. is supported by a National Health and Medical Research Council Investigator Grant (grant no. APP 2026403). A.Z. is supported by a Future Fellowship from the Australian Research Council (grant no. FT220100091).

BCP and HCP-D were each supported by the National Institute Of Mental Health of the National Institutes of Health under Award Number U01MH109589 and 1U01MH110274. Data were provided in part by the HCP, WUMinn Consortium (principal investigators D. Van Essen and K. Ugurbil; 1U54MH091657) funded by the 16 National Institutes of Health institutes and centres that support the National Institutes of Health Blueprint for Neuroscience Research; and by the McDonnell Center for Systems Neuroscience at Washington University. Additional data were provided by the PNC (principal investigators H. Hakonarson and R. Gur; phs000607.v1.p1).

Support for the collection of the PNC was provided by grant RC2MH089983 awarded to R. Gur and RC2MH089924 awarded to H. Hakonarson. This publication does not necessarily reflect the views of the funding agencies.

