## Supplementary Information for "Frontolimbic Functional Connectivity Development from Birth to Emerging Adulthood"

*Corresponding Authors:

Wonyoung Kim, M.A.

**This PDF file includes:**

Supporting text

Figures S1 to S19

Tables S1 to S11

Legends for Movies S1 to S4

SI References

**Other supporting materials for this manuscript include the following:**

Movies S1 to S4

**Contents**

**Supporting Methods**

**Supporting Figures**

Figure S1. GAMLSS Model Diagnostics

Figure S2. Developmental Pattern of Differentiation Index Across Amygdala and Hippocampus

Figure S3. Lateral Frontal Connectivity Maps Across Development

Figure S4. One-year Age Bin Maps of Hippocampus Medial Frontal Connectivity

Figure S5. One-year Age Bin Maps of Hippocampus Lateral Frontal Connectivity

Figure S6. One-year Age Bin Maps of Amygdala Medial Frontal Connectivity

Figure S7. One-year Age Bin Maps of Amygdala Lateral Frontal Connectivity

Figure S8. Fronto-hippocampus and Fronto-amygdala Connectivity Differentiation Following the Sensorimotor-Association Axis Hierarchy

Figure S9. Voxelwise Pattern of Frontolimbic Differentiation Development in the PNC

Figure S10. Medial Frontal Connectivity Maps Across Development in the PNC

Figure S11. Lateral Frontal Connectivity Maps Across Development in the PNC

Figure S12. Functional Implications of Frontolimbic Differentiation Deviation in the PNC

Figure S13. Normative Maturational Trajectory Using Regression-Based Differentiation Index

Figure S14. Reference Maps from Scan 1 and Scan 2

Figure S15. Reference Maps from Three Age-bins Across Young Adulthood

Figure S16. Age Trajectory of Frontolimbic Differentiation in BCP without Cross-Cohort Harmonization

Figure S17. Age Trajectory of Frontolimbic Differentiation in HCP-D without Cross-Cohort Harmonization

Figure S18. Correlation Pattern of Cumulative Adversity on Fronto-amygdala Connectivity

Figure S19. Correlation Pattern of Cognitive Ability on Fronto-hippocampus Connectivity

**Supporting Tables**

Table S1. Regions of Interest Definition

Table S2. GAMLSS Distribution Choice

Table S3. Threshold-Free Cluster Enhancement Results in BCP

Table S4. Threshold-Free Cluster Enhancement Results in HCP-D

Table S5. Spatial Permutation Test of BCP Subregion Significance

Table S6. Spatial Permutation Test of HCP-D Subregion Significance

Table S7. Multiple Regression Model Statistics for the Spatial Axes

Table S8. Item Importance of Adversity Association

Table S9. Domain Importance of Cognition Association

Table S10. Behavior Measures in the PNC

Table S11. Reference Map Similarity in Random Half-Splits

**Supporting Movies**

Movie S1. Hippocampus Connectivity Pattern Across Medial Frontal Cortex

Movie S2. Amygdala Connectivity Pattern Across Medial Frontal Cortex

Movie S3. Hippocampus Connectivity Pattern Across Lateral Frontal Cortex

Movie S4. Amygdala Connectivity Pattern Across Lateral Frontal Cortex

**Methods**

**Datasets and Participants**

We utilized resting-state scans from multiple large-scale datasets collected at different sites: the Human Connectome Project Young-Adult (HCP-YA) dataset, the Human Connectome Project - Development (HCP-D) dataset, and the Baby Connectome Project (BCP). The HCP-YA is the main dataset of the HCP initiative designed to map brain connectivity and its variability using advanced multimodal MRI (N = 1,200, ages 22-35) (1). The HCP-D is a part of the lifespan neuroimaging initiative of the HCP comprising 652 individuals aged 5-21 (2). Lastly, the BCP is an early life dataset with longitudinal follow-up scans that study brain development from birth to early childhood (N = 297, ages 0-5) (3).

The normative adult reference connectivity patterns were generated from the HCP-YA dataset (HCP-YA S1200 release). T1-weighted images were acquired with the following parameters: TR = 2,400 ms, TE = 2.14 ms, and resolution = 0.7 × 0.7 × 0.7 mm^3^. Resting-state fMRI images were acquired using the EPI sequence with the following parameters: TR = 720 ms, TE = 33.1 ms, resolution = 2 × 2 × 2 mm^3^, and total 4800 volumes.

The HCP-D dataset was used to cover the age range from late childhood to emerging adulthood (male/female: 301/351). T1-weighted images were acquired with the following parameters: TR = 2,500 ms, TE = 1.8 / 3.6 / 5.4 / 7.2 ms, and resolution = 0.8 × 0.8 × 0.8 mm^3^. Resting-state fMRI images were acquired using the EPI sequence with the following parameters: TR = 800 ms, TE = 37 ms, resolution = 2 × 2 × 2 mm^3^, and total 1952 volumes.

BCP was analyzed to cover connectivity development from birth to early childhood (male/female: 140/157). T1-weighted images were acquired with the following parameters: TR = 2,400 ms, TE = 2.24 ms, and resolution = 0.8 × 0.8 × 0.8 mm^3^. Resting-state fMRI images were acquired using the EPI sequence with the following parameters: TR = 800 ms, TE = 37 ms, resolution = 2 × 2 × 2 mm^3^, and 420 volumes.

**Neuroimaging Data Processing**

The HCP-D dataset is distributed minimally preprocessed with ICA-FIX denoising applied (4, 5). We additionally employed steps for data denoising and sanitation. First, scanning runs with > 20% of frames exceeding framewise displacement (FD) > 0.5 mm were excluded. Then, individuals with all four resting-state runs and a mean FD of < 0.2 mm were retained, resulting in the final sample size of *N* = 627 (337 females and 290 males) for the main analyses. To further alleviate the confounding effects of motion, we censored frames with FD > 0.5 mm and their adjacent frames (one before and two after), with linear interpolation replacing the missing frames to retain temporal continuity. For additional denoising and sanitation, the first four volumes were removed, 24 motion parameters were regressed out, and high-pass filtering was carried out. The residual timeseries data were normalized within run and concatenated across runs.

The HCP-YA followed the same pipeline to establish the adult reference fronto-amygdala and fronto-hippocampus connectivity maps from the final sample of *N* = 1,084. Of note, for the main analyses we utilized the two runs from the first resting-state scan, amounting to 2400 volumes. The second scan was processed in the identical pipeline to be used as a reproducibility check for the reference connectivity pattern (*N* = 1,031). The reference connectivity maps calculated from the second scan was nearly identical (Fig. S14; r = 0.991 for fronto-hippocampus connectivity map; r = 0.988 for fronto-amygdala connectivity map). As it was confirmed that the reference connectivity maps are reproducible over different days of scanning, we proceeded with maps derived from the first scan.

The BCP dataset was preprocessed using Nibabies v25.2.1. (6), the infant-equivalent of fMRIPrep, built atop Nipype 1.10.0 (7). To account for the distinct morphology of the developing brain, NiBabies adapts the standard fMRIPrep framework by substituting adult-centric components with age-appropriate alternatives, such as infant-specific templates and Infant FreeSurfer (8, 9). Individuals with missing structural or resting-state data, major neurological conditions, preprocessing failures, mean FD over 0.5 mm, or noticeable artifacts in the structural images were excluded. Runs with > 20% of frames exceeding framewise displacement (FD) > 0.5 mm were excluded as well. For individuals with multiple runs in a session, we used the run with the lowest movement determined by mean FD to ensure that all individual data points were comparable in scan length. Preprocessing steps included motion correction, alignment with structural images using boundary-based registration, censoring with interpolation, removal of first four frames, 24 motion parameters and white matter and cerebrospinal fluid signal regressed out, high-pass filtering, and projection onto CIFTI space. The residual data were normalized, with the final sample size of 195 individuals with the total of 412 scans. For the main cross-sectional analyses, we used the last scan of each individual to balance the data distribution skewed toward infancy. All scans were used for the sensitivity analyses for cohort effects. Although the exclusion based on movement was more lenient in the BCP compared to the HCP-D, this was still more stringent than recent work that successfully utilized the same dataset to map functional connectivity in infants (10). The additional denoising steps and adding mean FD as a covariate as in the main analysis ensured that motion was adequately controlled for in the ensuing analyses.

**Fronto-hippocampus and Fronto-amygdala Functional Connectivity Pattern Differentiation**

Amygdala and the hippocampus were identified using the Melbourne Subcortex Atlas (11), while the frontal cortex vertices were located using the Glasser atlas (12). The Melbourne Subcortex Atlas delineates the hierarchical organization of subcortical regions based on gradients of functional connectivity profiles with the entire cortex. The Glasser atlas defines regions of the cortex using cortical architecture, function, connectivity, and topography. We defined the frontal cortex to include the orbitofrontal cortex, medial frontal cortex, and the lateral frontal cortex anterior to the somatosensory regions that have cortical characteristics distinct from frontal cortex regions. Exact regions included are listed in Table S1.

The differentiation index (DI) was calculated for each voxel in an individual as follows:

$$DI=Corr\left( DevFC, RefFC_{amyg} \right)-Corr(DevFC,RefFC_{hipp})$$

where a developing individual’s frontal cortex normalized connectivity pattern is denoted *DevFC*. *RefFC_amyg_* and *RefFC_hipp_* indicate the adult reference connectivity patterns across the frontal cortex for either the amygdala or the hippocampus. In detail, for each participant in the developmental cohorts, frontolimbic functional connectivity patterns were calculated per voxel by deriving the correlation between the timeseries of a hippocampus or amygdala voxel and those of all frontal vertices. This was repeated for all voxels in the amygdala and the hippocampus, and all connectivity measures were subsequently normalized through Fisher’s Z transform. Then, we calculated two spatial correlations per each voxel: one measuring the similarity of their fronto-amygdala connectivity pattern to the reference adult fronto-amygdala connectivity pattern, and another for the hippocampus. The differentiation index was defined as the voxel-wise difference between these two correlation values. In essence, the differentiation index asks the question for each brain voxel: “does its connection patterns to the frontal cortex look more like a typical adult amygdala’s pattern or a typical adult hippocampus’ pattern?”. This differentiation index provides an intuitive and continuous measure of circuit identity differentiation in terms of frontal connectivity patterns. A positive differentiation index value in a seed voxel (e.g., within the amygdala) indicates that its frontal connectivity pattern is more similar to the adult amygdala pattern than to the adult hippocampus pattern, signifying a more differentiated state. As such, amygdala voxels are expected to have positive differentiation index values. On the other hand, negative differentiation index values indicate frontal connectivity patterns closer to the adult hippocampus pattern than the adult amygdala pattern. For conceptual clarity, differentiation indexes in the hippocampus were multiplied by -1 so that higher differentiation index in both the amygdala and the hippocampus track the degree of differentiation in the same direction. ComBAT harmonization was applied to account for site effects (13). Lastly, differentiation index was averaged for the amygdala and the hippocampus respectively for statistical analyses.

**Normative Development of Fronto-hippocampus and Fronto-amygdala Connectivity Differentiation**

First, Generalized Additive Model for Location, Scale, and Shape (GAMLSS) models were fit to accurately model the normative trajectory of the average connectivity differentiation for the amygdala and the hippocampus, respectively (14). Intracranial volume (ICV) and mean FD were regressed out of the amygdala and hippocampus average differentiation index prior to fitting. Age and sex were entered into the model with site also added as a random effect term. Race was omitted from the model as several unique categories of race led to failure in model convergence across cross-validation folds. We used fractional polynomials to estimate the nonlinear trends for its balance in flexibility in model fitting and simplicity in model choice (15). Fitting the GAMLSS models involved a combination of decision criteria. In detail, all possible distributions were tested, and the resulting models were compared in their Akaike Information Criterion (AIC). The final model parameters were selected based on a combination of criteria, including low AIC, successful convergence across all cross-validation folds, and satisfactory model fit as confirmed by residual plots, density estimate plots, normal Q-Q plots, and visual inspection of the fitted lines. Distribution choice and model diagnostics is described in Table S2 and Fig. S1. Finally, the z-scores that quantify the individual deviations from the estimated norm was calculated in a 10-fold cross-validation scheme for each of the two normative models. The z-scores were used in all subsequent statistical analyses to characterize individual deviations.

To isolate and compare the effect of age across the two circuits, we derived cumulative maturational curves (Fig. 2B). The curves characterized the developmental progress made at a given age for fronto-hippocampus and fronto-amygdala differentiation, respectively, in percentages. To this end, we extracted the non-linear age effects from the GAMLSS μ term, isolating the median value of differentiation at each age controlling for sex and site effects. For each circuit, this median value at the earliest age was subtracted from the median values across the entire age range before scaling to the maximum value of differentiation present in the sample. This in effect expressed the cumulative developmental progress made at each age. Since normalized to their respective maximum points, the developmental timings of fronto-hippocampus and fronto-amygdala differentiation could now be directly compared. As an illustration, the cumulative percentage at age of 1 year was calculated for both circuits.
**Connectivity Changes in Limbic Structures in Relation to Differentiation**

We then examined the effect of age on the voxel-wise differentiation index maps of the individuals to identify voxels that became increasingly differentiated with age in frontal connectivity patterns (see Fig.S2A), using *randomise* in FSL. Threshold-Free Cluster Enhancement was used to circumvent arbitrary thresholds in detecting significant voxel clusters (16). As non-linear age trajectories were identified in both amygdala differentiation and hippocampus differentiation, we carried out the analyses separately for the two developmental cohorts. This allowed us to identify regions with pronounced age effects in each of the two developmental stages. Results are outlined in Tables S3-4.

To test if the effects we observed were significantly localized to subregions of the amygdala or the hippocampus, we carried out spatial permutation tests. First, the subregions were defined using the Melbourne Subcortex Atlas that allows finer parcellation of subcortical structures based on functional connectivity (11). The amygdala was further parcellated into medial and lateral subregions, and the hippocampus was divided into anterior and posterior subregions. Here, we summed the t values of the age effects for a subregion. If a subregion had higher summary t value than the other, we treated this as evidence of discrete subregion-specific effect. Next, BrainSMASH was conducted to make spatially permuted null t maps across the amygdala or the hippocampus that preserves the spatial autocorrelation (17). In each of the null maps, t values were again summed for the subregion. If the age effects were spatially constrained to any subregion, the true t value sum would be significantly greater than the majority of the permuted t value sums. This procedure was repeated for all amygdala and hippocampus subregions. Results are detailed in Tables S5-6.

Finally, to test the continuous effects of age across the voxel-wise differentiation index values that subregion analyses may have not fully captured, spatial axes analyses were performed. In detail, we tested multiple regression with the t value as the dependent variable and coordinates in MNI space (mm) along the X axis (medial-lateral axis), Y axis (posterior-anterior axis), and Z axis (dorsal-ventral axis) as independent variables. Absolute values were used for the X axis so that its values would indicate the medial-to-lateral direction for both the left and right hemispheres. If the t values change as functions of spatial axes, we treated this as evidence of continuous subregion-specific effect. This identified which spatial axis the developmental differentiation effects significantly changed along. Three multiple regression models were tested: 1) a model including all hippocampus and amygdala voxels, 2) a hippocampus model that highlights within-hippocampus gradients, and 3) an amygdala model that shows within-amygdala gradients. For visualization, we projected the predicted values of the multiple regression models back onto the voxel coordinates (Fig. S2B).
**Connectivity Changes in Frontal Cortex in Relation to Differentiation**

To characterize the frontal cortex connectivity patterns varying across age and across individuals, we created group-level maps of frontolimbic connectivity patterns stratified by age and differentiation deviation (Fig. 3 and Fig. S3). We chose four group levels for age to visualize the contrasting effects most effectively, with each of the younger and the older cohort divided at the median age (age = 1 for the birth to early childhood cohort; age = 14 for the late childhood to emerging adulthood cohort). Visualization with single year age bins is also available in Fig. S4-7. The reference of comparison was the adult amygdala-specific or hippocampus-specific frontal connectivity map, created through computing the difference between the reference fronto-amygdala and fronto-hippocampus connectivity maps from adults. Spatial correlation between the comparison reference maps and group-level maps were charted as a descriptive depiction, which in nature largely recapitulates the individual-level differentiation trend across age.

Next, the frontal cortex connectivity patterns were compared to known cortical organization schemes to contextualize the connectivity shift beyond visual descriptions. Comparison with the 7 canonical functional networks (18) was motivated by previous literature that amygdala and hippocampus, in healthy adults, are each uniquely connected to the SN and the DMN (19, 20). As it was visually evident that the networks on their own were not sufficient to describe the connectivity shifts in full, we utilized the S-A axis that explains cortical organizations, including the functional networks, in a continuous expression of hierarchy (21). In detail, in the one-year age bin group-level maps of the fronto-hippocampus connectivity and fronto-amygdala connectivity, we averaged the normalized connectivity values across the frontal cortex for each network. This resulted in mean frontal connectivity across 5 networks (DMN, Control Network, Limbic Network, SN, and the Dorsal Attention Network) and across 22 age band maps for both the amygdala and the hippocampus. Plotting this across age showed clear age trends involving all functional networks of the frontal cortex rather than only SN and DMN. We next built a linear model as follows:

$$FrontConn=\beta_{0}+\beta_{1}Age+\beta_{2}SAhier+\beta_{3}\left( Age\times SAhier \right)+\beta_{4}Circ+\beta_{5}\left( Age\times Circ \right)+\beta_{6}\left( SAhier\times Circ \right)+\beta_{7}\left( Age\times SAhier\times Circ \right)+\epsilon$$

$$\mathrm{where}$$

*FrontConn*, *SAhier*, and *Circ* indicate mean frontal cortex connectivity within a functional network, the network’s mean position in the S-A hierarchy, and circuit (i.e., fronto-amygdala or fronto-hippocampus). The core aim was to test the three-way interaction between the network average S-A hierarchy ranking, age, and circuit in predicting the network average connectivity. As fronto-hippocampus was assigned 0 and fronto-amygdala 1 in the *Circ* term, $\beta_{3}$ expressed in the fronto-hippocampus connectivity how networks higher and lower in S-A hierarchy may have different age trends (e.g., increase or decrease). Conversely, $\beta_{3}+\beta_{7}$ expressed the same in the fronto-amygdala connectivity. Same modeling was repeated with the SN, the DMN, or both excluded to ensure that the effect was driven by the overall S-A hierarchy across the frontal cortex, not only the SN and the DMN (Fig. S8). To directly compare the amygdala and the hippocampus within this framework, we also calculated per each network the age slope of amygdala – hippocampus connectivity strength that varies as a function of the S-A hierarchy position. In specific, across all one-year age bin connectivity maps, network-average hippocampus connectivity strength was subtracted from network-average amygdala connectivity strength, then correlated with age.

**Association with Adversity and Cognitive Ability**

We tested the hypothesis that the cumulative adverse life events in the past year were associated with the differentiation z-scores (i.e., deviation from the age-specific median). Cumulative adversity was calculated as the sum of adverse life events experienced in the past year, as reported in a questionnaire as a part of the HCP-D data collection (22). We also explicitly tested for the interaction term of sex, age, and socioeconomic status in separate models given reports of divergent maturational trajectories between males and females (23), age-specific effects of adversity (24), and influence of socioeconomic status on brain development (25). To account for the possibility that the association between the differentiation z-scores and cumulative adversity is driven by how socioeconomic status, recent stress, or perinatal stress exposure may influence fronto-amygdala or fronto-hippocampus connectivity development (26-30), additional partial correlations were conducted controlling for such variables.

Next, in order to scrutinize the contributions of the adverse life events, we built a Principal Component Regression (PCR) model that uses the 25 adverse life event items to predict the amygdala differentiation z-score. Bootstrapping was carried out 10,000 times to describe whether a feature was important within the PCR models, robust to potential sampling biases. Feature importance was calculated by aggregating the principal component loadings across the items that together predicted the differentiation z-score (mean r_predicted-actual_ = 0.133, 95% CI [0.02 0.23]). The significance for each item was assessed by examining the confidence intervals calculated through bootstrapping. If the 95% confidence interval of the principal component loading for an item did not contain 0, the item was deemed to be statistically significant, equivalent to a p-value being smaller than 0.05. Correction for multiple comparisons was performed using the False Discovery Rate (31).

We next tested the hypothesis that individual deviations in differentiation were related to individual differences in cognitive ability at a given age. Cognitive ability was measured through a composite of cognitive assessments in the NIH toolbox (32). This includes Dimensional Change Card Sort Test, Flanker Inhibitory Control and Attention Test, List Sorting Working Memory Test, Oral Reading Recognition Test, Pattern Comparison Processing Speed Test, Picture Vocabulary Test, and Picture Sequence Memory Test, each designed to tap into distinct domains of intelligence. The measures were summed and age-adjusted to yield the total cognitive ability score at a given age. Interaction of sex, age, and socioeconomic status were also tested (33, 34). Again, partial correlations controlling for socioeconomic status, perinatal exposure to stress, and recent stress exposure were conducted to rule out the possibility of the effects being driven by how these factors influence connectivity development.

We applied the same feature importance analysis to the association between the hippocampus differentiation z-score and cognitive ability. PCR model that uses 7 age-adjusted cognitive test scores to predict hippocampus differentiation z-score was used to understand the contributions of each cognitive test (mean r_predicted-actual_ = 0.135, 95% CI [0.02 0.25]). Bootstrapping with 10,000 iterations provided confidence intervals for both the PCR model and the feature importance. Correction for multiple comparisons was performed using the False Discovery Rate (31).
**Replication in a Community Dataset**

We sought to replicate our key findings in a community sample, the Philadelphia Neurodevelopmental Cohort (PNC). The PNC is a large-scale community dataset that aims to capture brain maturation and risk for mental illness (*N* = 1,395, ages 8-23) (35). PNC subjects were recruited across the Children’s Hospital of Philadelphia network where they received medical care outside of psychiatric services. Some individuals involved in the PNC cohort have clinical and sub-clinical psychiatric symptoms (36), thus capturing a wider range of the population spectrum than the main datasets.

For the PNC, T1-weighted images were acquired with the following parameters: TR = 1,810 ms, TE = 3.5 ms, and resolution = 1.0 × 1.0 × 1.0 mm^3^. Resting-state fMRI images were acquired using the EPI sequence with the following parameters: TR = 3000 ms, TE = 32 ms, resolution = 3 × 3 × 3 mm^3^, and total 124 volumes.

The PNC dataset was preprocessed using fMRIPrep version 25.1.1 (37) based on Nipype 1.10.0 (7). Individuals with missing structural or resting-state data, major neurological conditions, preprocessing failures, mean FD over 0.2 mm, or noticeable artifacts in the structural images were excluded. The same denoising procedures as done in the HCP-D dataset were done. This included censoring with interpolation, removal of first four frames, 24 motion parameters and white matter and cerebrospinal fluid signal regressed out, and high-pass filtering. The residual data were normalized, with the final sample size of N = 898.

The same set of analyses from the main findings were carried out in the PNC dataset, from estimating normative trajectory of differentiation to characterizing differentiation patterns. To test for replication, we compared between the HCP-D dataset and the PNC dataset the voxel-wise maps of age effects on differentiation index across the amygdala and the hippocampus. We expected to find positive differentiation index values in the amygdala voxels and negative differentiation index values in the hippocampus as confirmed in the HCP-D voxel-wise map (see Fig. S2A and Fig. S9). The effect of differentiation was also inspected across the frontal cortex vertices as done before (Fig. S10-11).

Functional implications were also investigated in the PNC although measurements differed from HCP-D (Fig. S12). As a proxy measure of adversity, we utilized the Kiddie Schedule for Affective Disorders and Schizophrenia (K-SADS) (38) Post-Traumatic Stress Disorder module items, which measured lifetime exposure to severe trauma events (e.g., “Were you ever in a serious fire?” or “When your parents got mad at you, did they hit you?”). As the nature of the adversity measured here is inherently different from the adverse life events questionnaire from the main analysis, this was also deemed an opportunity to test if the main findings can be extended into adversity that occurred beyond a year past and that are more severe. Cognitive ability score was calculated as a summary score of cognitive test performance metrics (e.g., True positive and false positive rates for Penn Face Memory Test). Scores that indicate higher intelligence at lower scores were inverted before standardizing all scores and summing to make a composite score. The full extent of items used for both the traumatic event exposure score and cognitive ability score are listed in Table S8.
**Sensitivity Analyses**
1. Regression-based differentiation metric

Although relatively intuitive and simplistic, our differentiation index does not formally address the high similarity between the adult amygdala connectivity and hippocampus connectivity across the frontal cortex (spatial r = 0.742). This may introduce concerns that for some voxels the discrepancy between resemblance towards the amygdala reference connectivity and the hippocampus reference connectivity for a developing individual’s voxel may be so small that it is indistinguishable from noise. To address such concerns, we constructed the differentiation index using an alternative method that utilizes linear regression models.

Specifically, a developing individual’s voxel connectivity pattern is defined as follows:

$$DevFC=1+beta_{1}* RefFC_{amyg}+beta_{2}* RefFC_{hipp}+\epsilon$$

where *DevFC* is a developing individual’s frontal cortex connectivity pattern and *RefFC_amyg_* and *RefFC_hipp_* are frontal cortex connectivity patterns of amygdala and hippocampus in adults. This linear model, fitted for each voxel, models resemblance toward the reference amygdala connectivity and the reference hippocampus connectivity simultaneously.

These are subsequently used to derive the regression-based differentiation index:

$$DI_{reg}=\frac{beta_{1}-beta_{2}}{\sqrt{(var\left( beta_{1} \right)+var\left( beta_{2} \right)-2cov\left( beta_{1},beta_{2} \right)}}$$

which is equivalent to a t-statistic, allowing statistical significance testing for how each voxel is differentiated.

Finally, a structure-average differentiation index for the amygdala is defined as follows:

$$DI_{reg}Amyg=\frac{\sum(DI{}_{reg}*amyg_{s})}{\sum amyg_{s}}-\frac{\sum(DI{}_{reg}*hipp_{s})}{\sum hipp_{s}}$$

where the subscript *s* marks voxels passing the significance testing (i.e., p < 0.05). *amyg_s_* signify a binary index indicating which amygdala voxels are significantly differentiated toward the amygdala reference, whereas *hipp_s_* is a binary index indicating which amygdala voxels are significantly differentiated toward the hippocampus reference. In other words, the average amygdala differentiation is calculated by summing the differentiation index of voxels significantly resembling the amygdala reference connectivity over the hippocampus’, divided by the number of significant voxels, which is then subtracted by the sum of differentiation index of voxels significantly resembling the hippocampus reference connectivity over the amygdala’s, divided by the number of significant voxels. If the amygdala as a whole approaches perfect differentiation, *amyg_s_* is close to the number of amygdala voxels where *hipp_s_* approaches 0, resulting in the *DI_reg_Amyg* becoming comparable to the simple sum of *DI_reg_* of all amygdala voxels. The structure-average differentiation index for the hippocampus is calculated using the same logic across hippocampus voxels. In effect, this regression-based differentiation index would be equivalent to the original differentiation index if all voxels that almost equally resemble the reference amygdala and the reference hippocampus were left out from the metric.

Normative trajectories of fronto-hippocampus and fronto-amygdala differentiation over development using this alternative differentiation metric are described in Fig. S13.
2. Stability of the Reference Connectivity Map

Our main analytic pipeline depends on spatial correlation of developing connectivity maps with the reference maps derived from adults. While offering simplicity in delineating a concerted developmental effect from highly multivariate feature sets (i.e., connectivity maps across 15,616 vertices), this requires the assumption that the reference maps are stable representations of adult frontolimbic connectivity. To test this assumption, we carried out three tests: stability of the reference maps estimated across different days, across 100 random half-split groups within the first day, and across three age-bins within young adulthood. Results are outlined in Fig. S14-15 and Table S9.
3. Cohort Effects

One challenge of building normative growth models with different cohorts is ensuring that the developmental effects observed are not artifacts of merging heterogeneous cohorts. To test this possibility, we separately mapped the developmental trends for the birth to early childhood cohort and the late childhood to emerging adulthood cohort separately. Within-cohort site effects were still removed through ComBAT harmonization (13).

First, we tested if the developmental timings and the developmental trajectories of frontolimbic connectivity differentiation were preserved when we model the early childhood cohort separately. Namely, we checked for the early increase in fronto-hippocampus differentiation after birth and no increase in fronto-amygdala differentiation. To this end, we leveraged the full dataset including the longitudinal follow-up data by applying linear mixed effects models to account for repeated measures, an approach that was not feasible when merging cohorts and modeling with GAMLSS. The results are shown in Fig. S16. Importantly, scanning conditions differed in the BCP dataset where children younger than the age of 3 years were scanned during natural sleep while children older than the age of 3 years had the option to be scanned either during sleep or awake conditions (3). To check for scanning condition effects, we additionally carried out a linear mixed effects modeling with longitudinal data in the BCP dataset as done previously in Fig. S16 but including only sleep-condition data. We focused on the sleep-condition data as the awake-condition data did not have a sample size sufficient for its own analysis (13 scans). Comparable to the original trajectories seen in Fig. S16, fronto-hippocampus differentiation, but not fronto-amygdala differentiation, developed during early childhood when restricting the analysis to sleep-condition scans (399 scans) (β=0.111, p=0.021 for fronto-hippocampus; β=-0.040, p=0.421 for fronto-amygdala differentiation). This suggests that the developmental trend of fronto-hippocampus and fronto-amygdala differentiation is not driven by the scanning conditions.

The developmental effects were also checked in the late childhood to emerging adulthood cohort when analyzed separately. We expected the normative trajectories for fronto-amygdala connectivity differentiation increasing throughout adolescence from near zero at late childhood, which is in line with the lack of differentiation from birth to early childhood. For fronto-hippocampus connectivity, we expected differentiation to show a stable line forming at a positive value, also in line with the early differentiation from birth to early childhood. The results are shown in Fig. S17.
4. Potential Confounds in Functional Links

Throughout development, the brain is exposed to a number of extrinsic factors that are known to impact connectivity maturation, including perinatal exposure to stress, the state of stress at the time of measurement, and socioeconomic status (26-30). To account for possibilities where the amygdala or hippocampus differentiation deviation is better explained by such factors, a Perinatal Risk Score, the distress score from the Perceived Stress Scale (39), and socioeconomic status were tested alongside cumulative adversity and cognitive ability for relationships with differentiation deviation.

The Perinatal Risk Score comprised of the count of pregnancy complications, substance use during pregnancy, and birth outcomes from developmental history items (33, 40-42). These factors together have been shown to confer risks of high allostatic load and poorer health later on in life (43). In detail, the substance use items included whether the birth mother consumed or used alcohol, tobacco, cocaine, marijuana, or heroin during pregnancy. Pregnancy complications included high blood pressure, preeclampsia, eclampsia, reports of heavy bleeding, pregnancy-related diabetes, anemia, previa, abruptio, or other problems with the placenta during pregnancy. Birth outcomes included premature birth, birth weight, cyanosis, slow heartbeat, no breathing, convulsions, or requiring oxygen at birth. With the exception of birth weight which was dichotomized to low (<2,500 g) or normal (≥2,500 g) to be assigned risk or no risk (44), all factors were already binary (i.e., “yes” or “no”). These items were summed to devise the Perinatal Risk Score.

Perceived stress is measured through 10 self-report questionnaire items that gauge stress levels in the past month, which can be a proxy measure for the current or very recent state of stress. For example, the items included “In the last month, how often have you been upset because of something that happened unexpectedly?” or “In the last month, how often have you felt that you were unable to control the important things in your life?”.

Socioeconomic status score was calculated using the demographic information provided at the time of data collection (2). Parental education and family income were each standardized and combined to make a composite socioeconomic status score. Any individual missing in either were omitted for the analyses.
5. Correlation Pattern for Functional Implication

Although our main analyses revealed that individual deviation in fronto-hippocampus and fronto-amygdala differentiation were associated with adversity and cognitive abilities, it is still possible that such functional links are more related to local connectivity changes that our differentiation index is capturing rather than the phenomenon of differentiation itself. To address this possibility, we first mapped out how each fronto-amygdala and fronto-hippocampus connectivity values are correlated with adversity and cognitive adversity, then compared such correlation patterns with the adult reference map of amygdala-specific (or hippocampus-specific) connectivity. If such correlation patterns are indeed similar to the adult maps of differentiated connectivity states, this would serve as converging evidence that the behavioral links we found are more related to the widespread pattern of connectivity differentiation rather than focal connectivity correlates. The results are shown in Fig. S18-19.

**Figures**

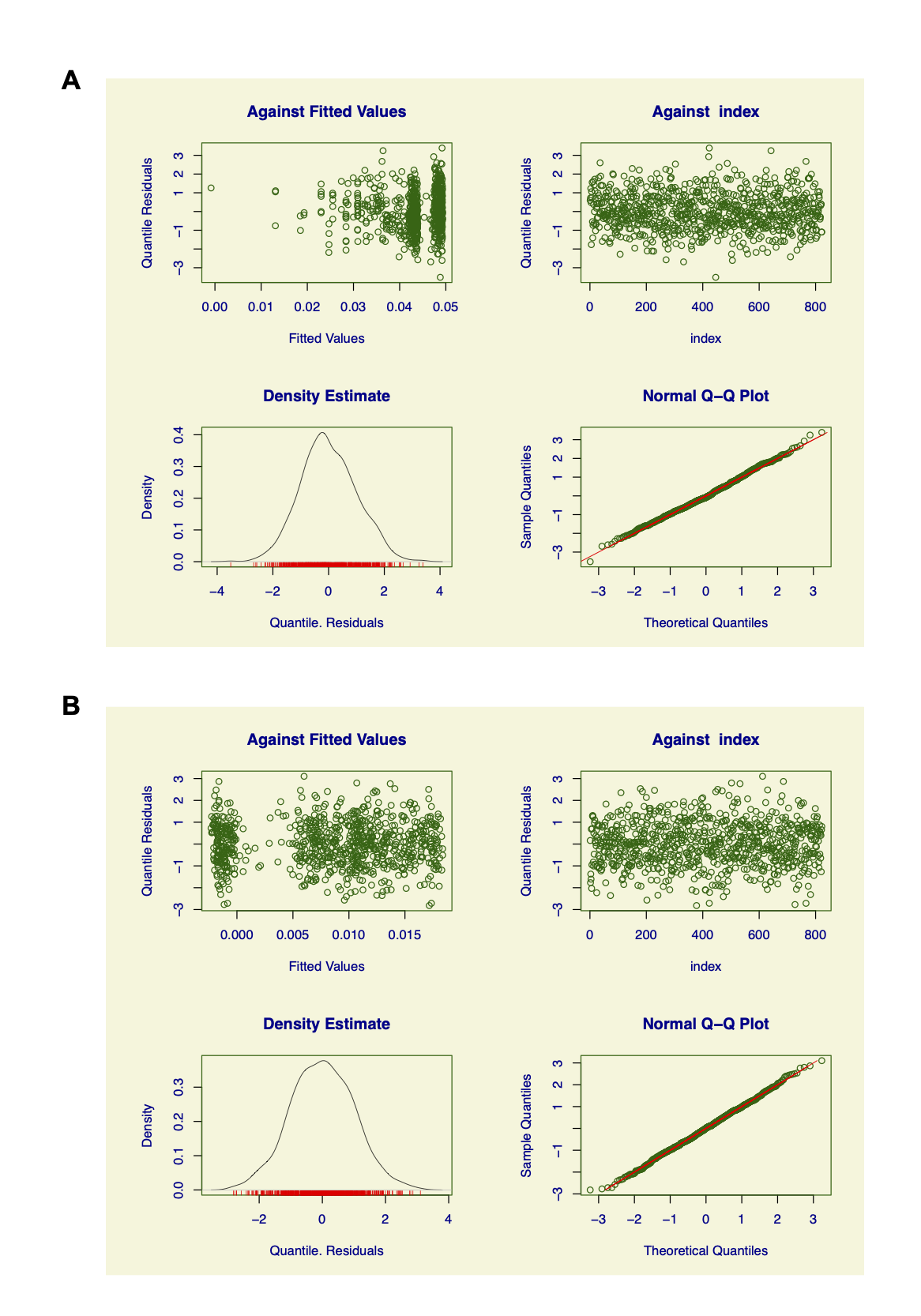

Figure S1. GAMLSS Model Diagnostics. **A** Model diagnostic information is shown for the fronto-hippocampus differentiation normative model. Upper panels indicate the quantile residuals against fitted values and against index, which is a check for homoscedasticity and independence of residuals. Lower panels show density estimate of quantile residuals as well as the normal q-q plot that check for normality. **B** Same as **A**, model diagnostic information is shown for the fronto-amygdala differentiation normative model.

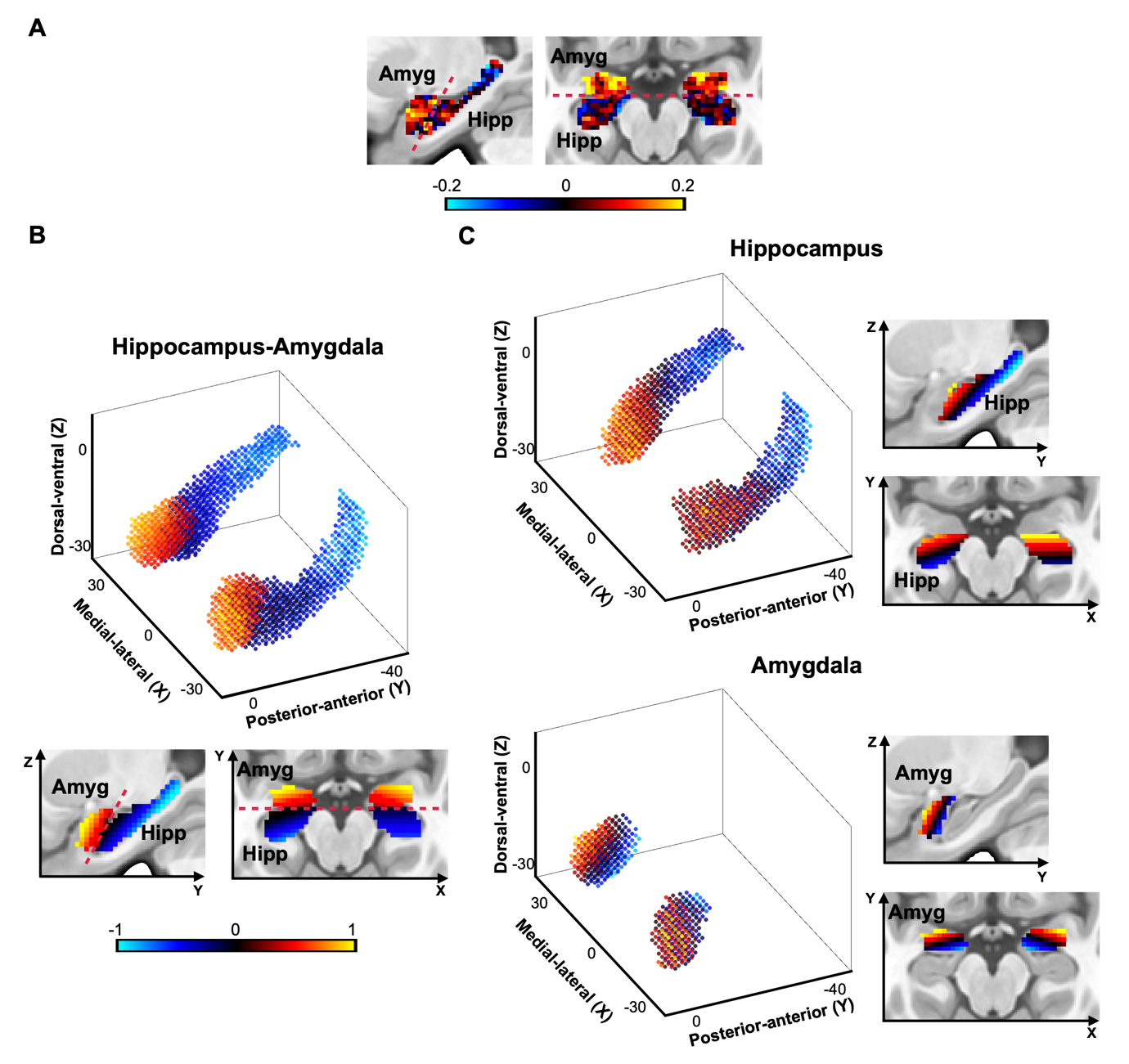

Figure S2. Developmental Pattern of Differentiation Index Across Hippocampus and Amygdala. **A** Age effects of voxel-wise differentiation index values. Positive values indicate a voxel’s connectivity pattern more strongly resembling the adult fronto-amygdala connectivity reference map more than the adult fronto-hippocampus connectivity reference map over age. Negative values indicate the converse. **B** Projected 3-dimensional model of differentiation index across hippocampus and amygdala. Values at each coordinate indicate the differentiation index value predicted by multiple regression models with the spatial coordinates as independent variables, standardized for each model to visualize the gradients clearly. At the global level, hippocampus and amygdala are clearly demarcated at their borders for connectivity differentiation. Panels below show 2-dimensional representations of the same model in anatomical space, similar to **A**. **C** Projected 3-dimensional models of differentiation index for hippocampus and amygdala, respectively. Developmental differentiation shows gradients of change within hippocampus and amygdala along the posterior-anterior (*Y*) axis and the medial-lateral axis (*X*). Panels to the right show 2-dimensional slices of hippocampus and amygdala respectively. The 2-dimensional representations are shown on a sagittal slice (*X* = 25.4mm) and an axial slice (*Z* = -18mm) in MNI space.

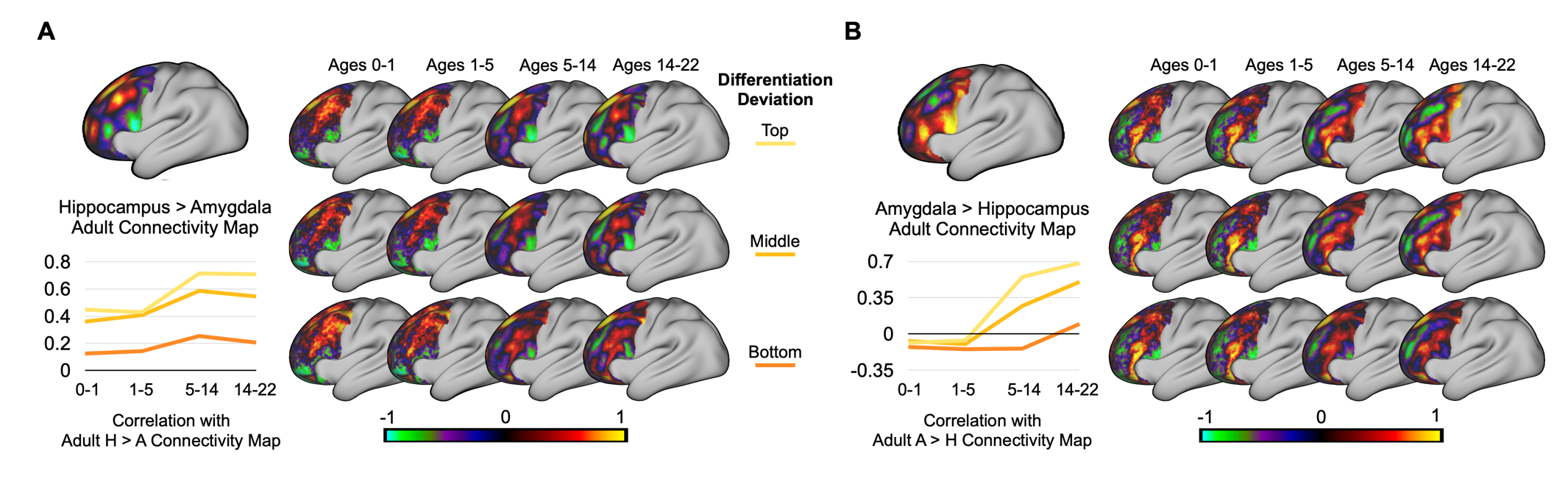

Figure S3. Lateral Frontal Connectivity Maps Across Development. **A** Fronto-hippocampus connectivity pattern shifts across ages and individual deviations in differentiation are shown. As the connectivity patterns become more differentiated, the negative connectivity in the ventral frontal cortex encroaches upwards to cover the dorsal lateral frontal cortex and the supplementary motor area. **B** Same as **A**, fronto-amygdala connectivity is shown stratified by age and differentiation deviation. Higher differentiation demonstrates the negative connectivity in the dorsal lateral frontal cortex diverging to form positive connectivity in the dorsal area, resembling the adult amygdala-specific connectivity map.

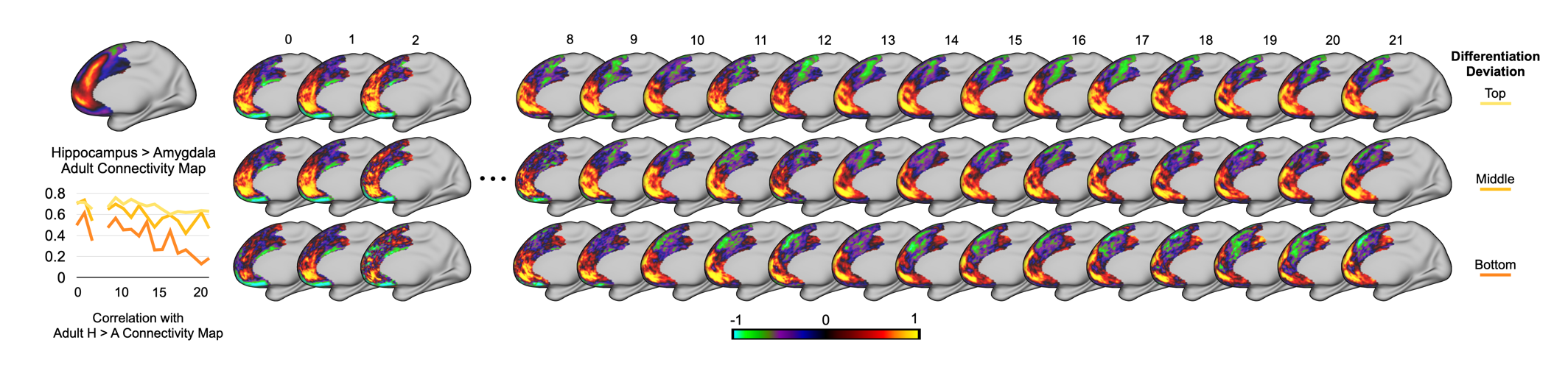

Figure S4. One-year Age Bin Maps of Hippocampus Medial Frontal Connectivity. Fronto-hippocampus connectivity patterns are shown for the medial frontal cortex in age bands of one year. Numeric values above each column of the connectivity maps mark the age bin that the maps correspond to.

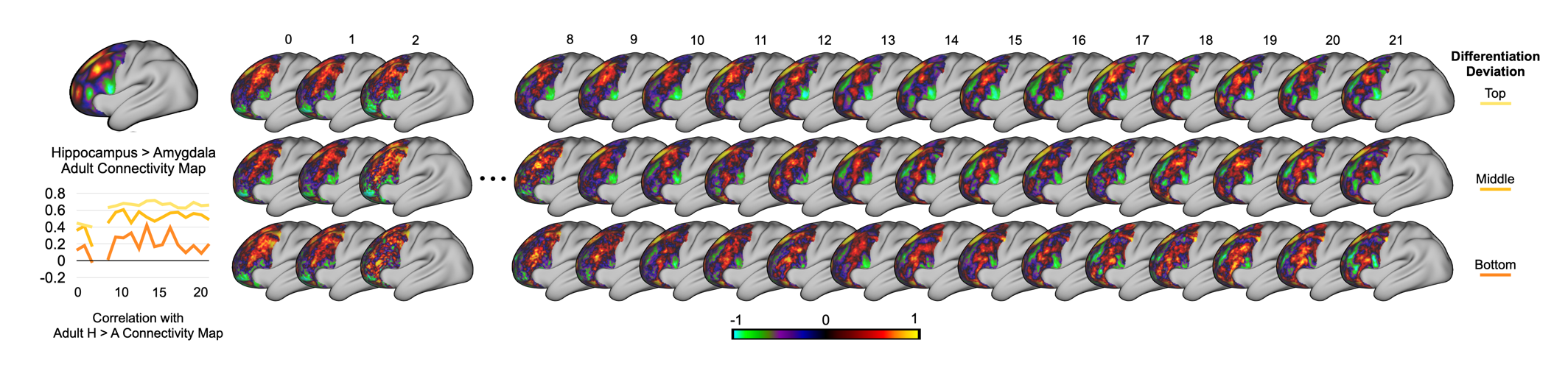

Figure S5. One-year Age Bin Maps of Hippocampus Lateral Frontal Connectivity. Fronto-hippocampus connectivity patterns are shown for the lateral frontal cortex in age bands of one year.

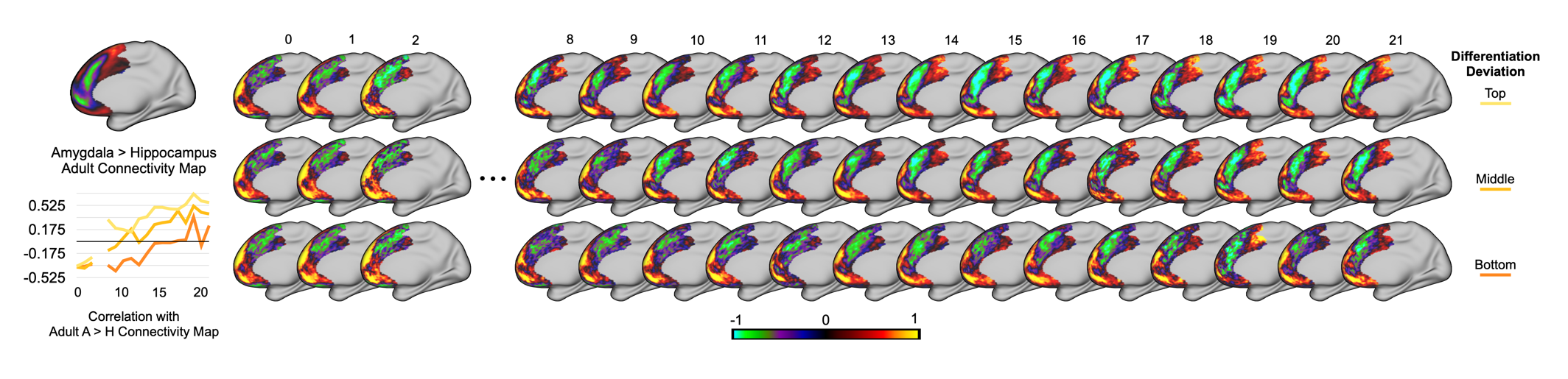

Figure S6. One-year Age Bin Maps of Amygdala Medial Frontal Connectivity. Fronto-amygdala connectivity patterns are shown for the medial frontal cortex in age bands of one year.

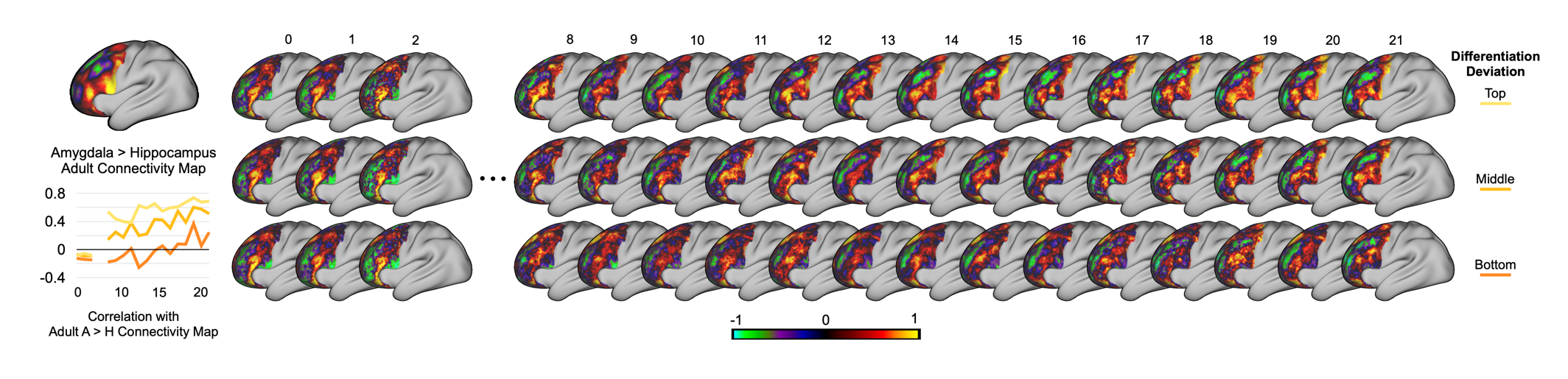

Figure S7. One-year Age Bin Maps of Amygdala Lateral Frontal Connectivity. Fronto-amygdala connectivity patterns are shown for the lateral frontal cortex in age bands of one year.

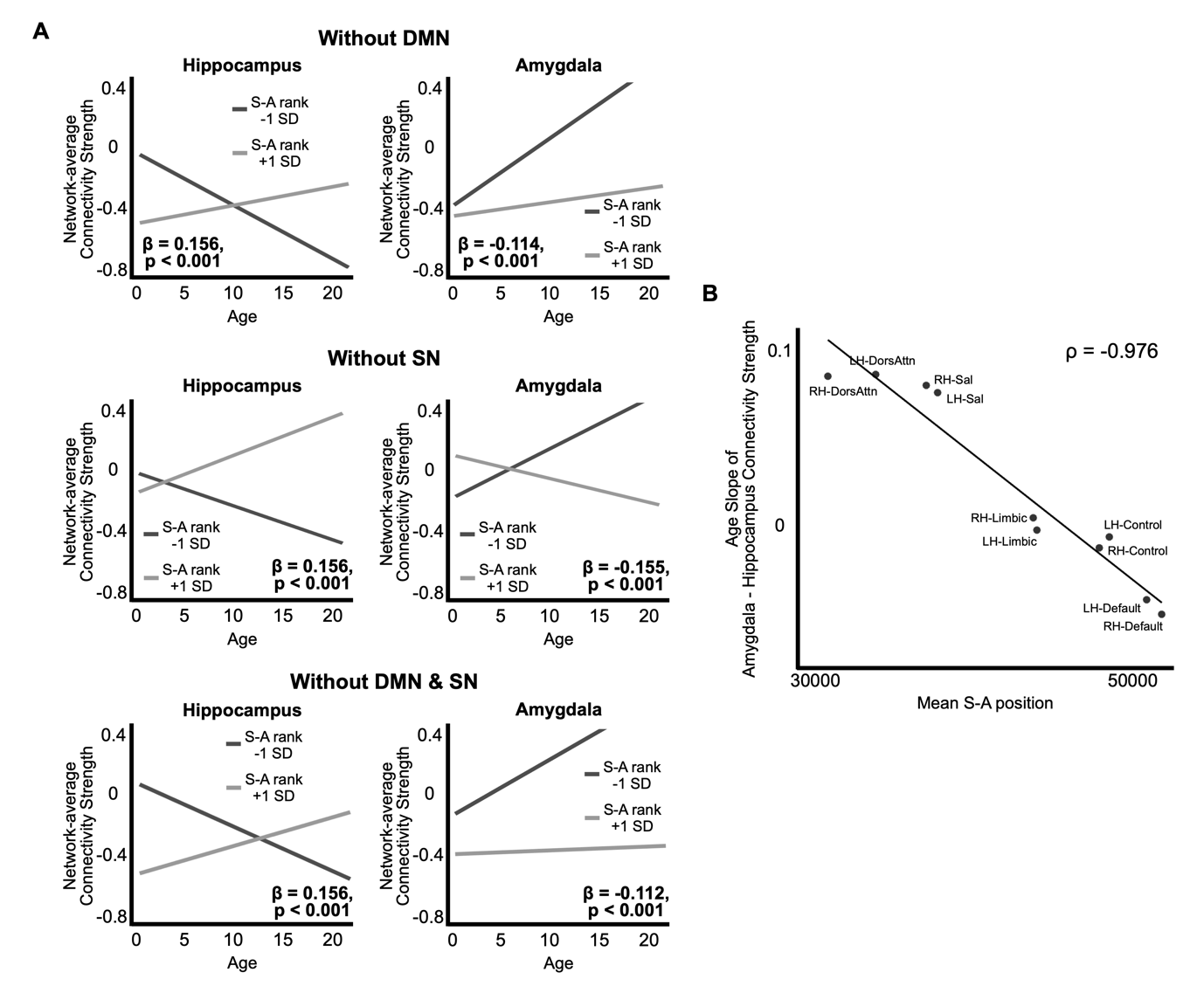

Figure S8. Fronto-hippocampus and Fronto-amygdala Connectivity Differentiation Following the Sensorimotor-Association Axis Hierarchy. **A** The same interaction analyses from the main findings were carried out without DMN, SN, or both. The interaction was significant across all exclusions, implying that the S-A hierarchy organization of fronto-hippocampus and fronto-amygdala differentiation was not driven by the simplistic amygdala-SN and hippocampus-DMN framework. **B** Frontal cortex network regions with lower mean S-A hierarchy position had higher age slope of amygdala – hippocampus connectivity strength, meaning connectivity with the amygdala increased over maturation when compared to connectivity with hippocampus. At the opposite end, network regions with higher mean S-A hierarchy position showed lower age slope, indicating hippocampus connectivity increasing over maturation compared to amygdala connectivity.

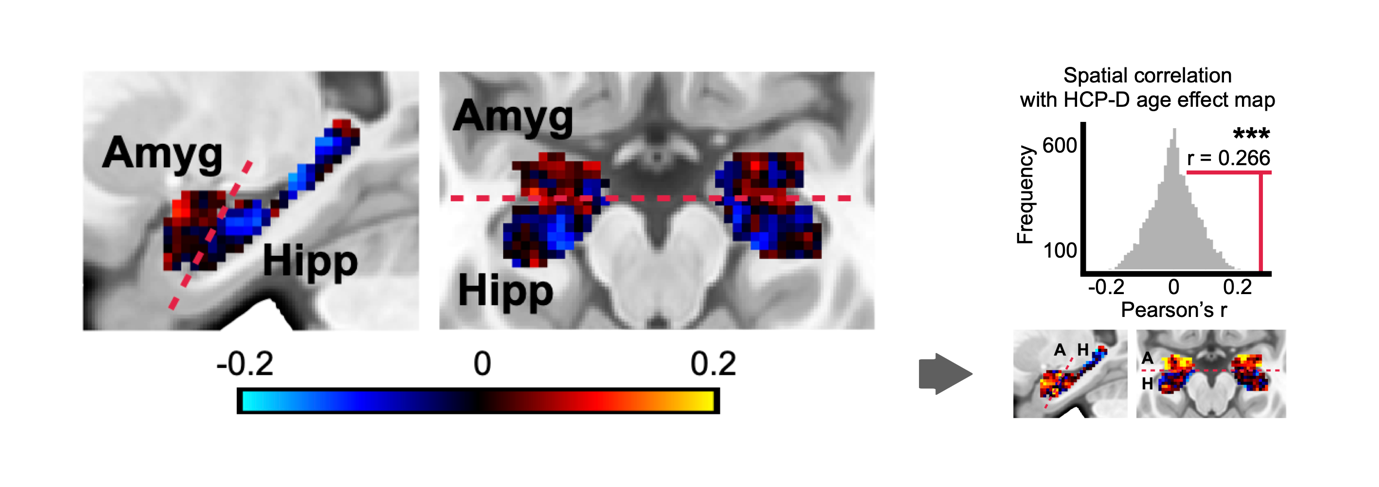

Figure S9. Voxelwise Pattern of Frontolimbic Differentiation Development in the PNC. Differentiation index is expressed across all voxels across the amygdala and the hippocampus. Same as Supplementary Figure 2A, positive values indicate differentiation towards the fronto-amygdala reference map and negative values indicate differentiation towards the fronto-hippocampus reference map. The pattern of voxels in the PNC is shown to be significantly similar to the one from HCP-D after BrainSMASH permutation, where amygdala voxels predominantly show positive values and hippocampus voxels show negative values, albeit with smaller effect sizes.

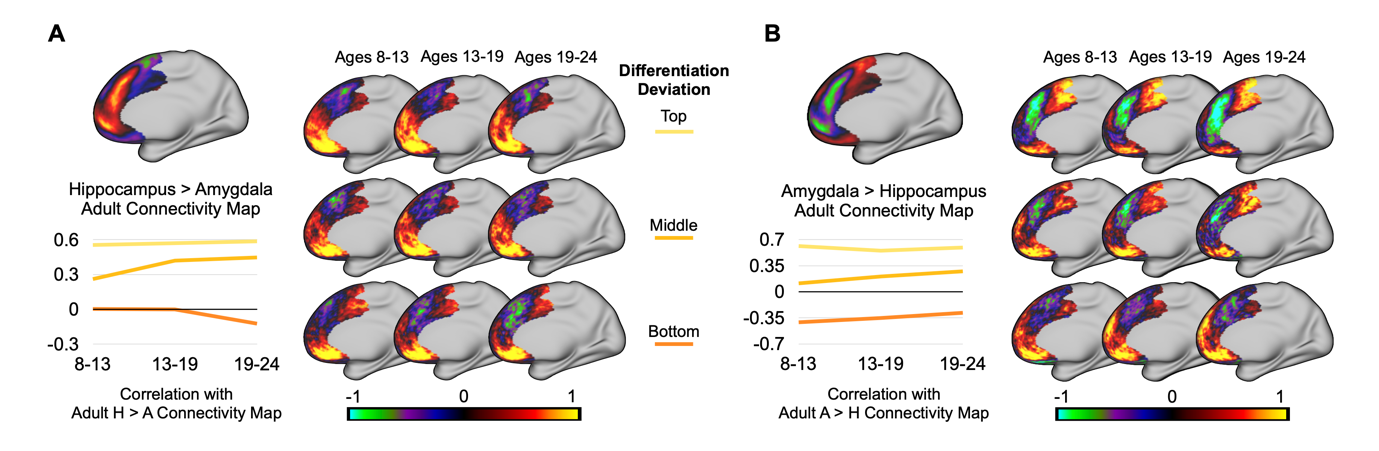

Figure S10. Medial Frontal Connectivity Maps Across Development

in the PNC. **A** Fronto-hippocampus connectivity maps are shown, stratified by age and differentiation deviation, both in tertiles. Though higher in individual variation, the group-level maps converge in that higher differentiation is marked by the patch of negative connectivity located in dorsal and posterior parts. **B** Same as **A**, fronto-amygdala connectivity maps are shown across age and differentiation deviation. Here, more differentiated connectivity maps demonstrate the patch of negative connectivity located to the ventral and anterior parts of the medial frontal cortex, mirroring the patterns from the main datasets.

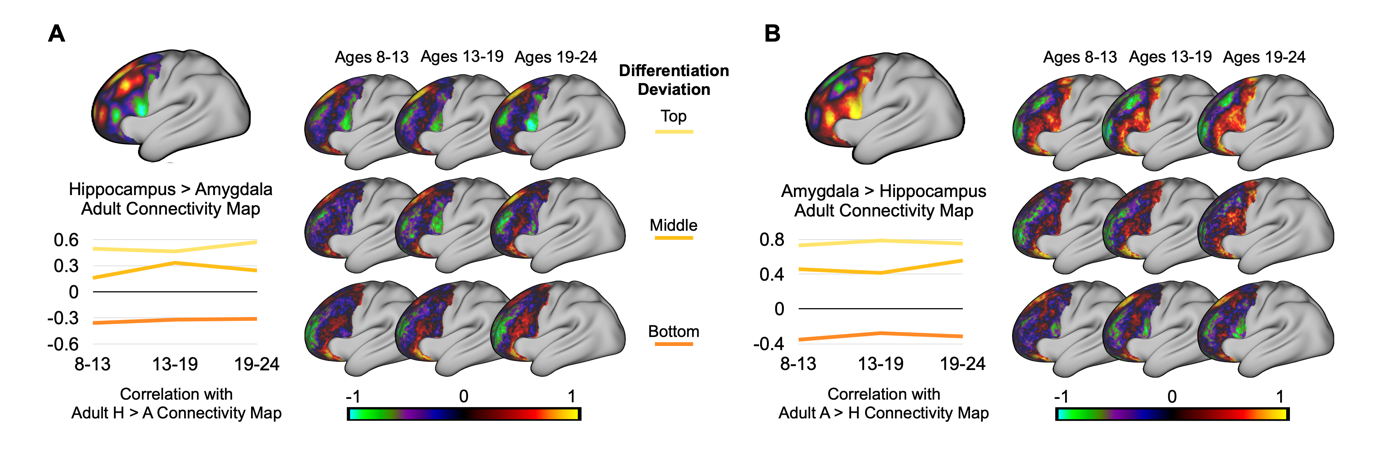

Figure S11. Lateral Frontal Connectivity Maps Across Development

in the PNC. **A** Fronto-hippocampus connectivity maps are shown, stratified by age and differentiation deviation, both in tertiles. Higher differentiation shows a negative patch of connectivity at the ventral part of lateral frontal cortex encroaching dorsally. **B** Same as **A**, fronto-amygdala connectivity maps are shown across age and differentiation deviation. As fronto-amygdala connectivity becomes more differentiated, the negative patch of connectivity spanning the dorsal and ventral parts splits into multiple patches.

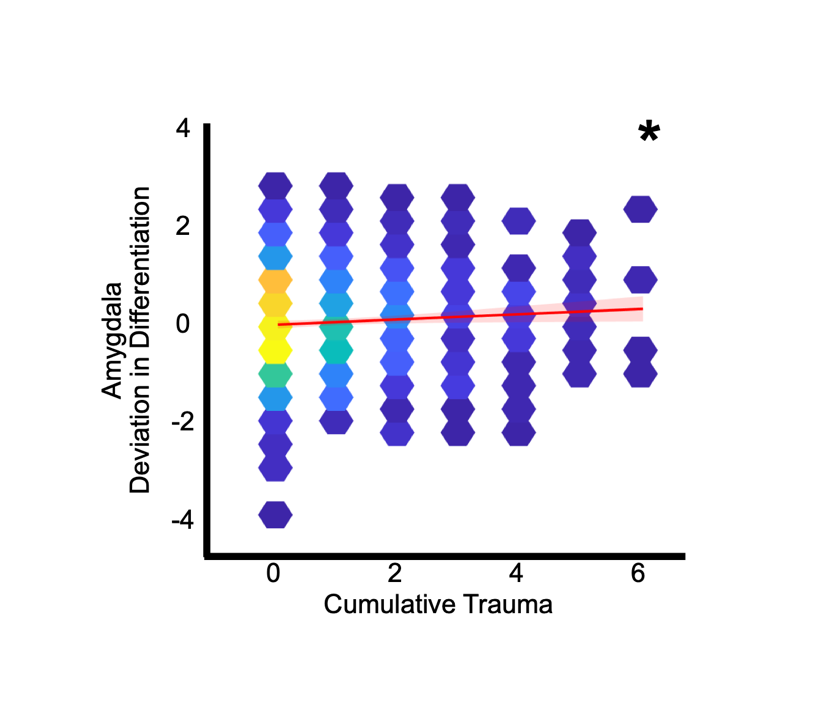

Figure S12. Functional Implications of Frontolimbic Differentiation Deviation in the PNC. Cumulative trauma measured from K-SADS PTSD module is positively associated with individual deviation in the amygdala differentiation at a given age (i.e., Z-score).

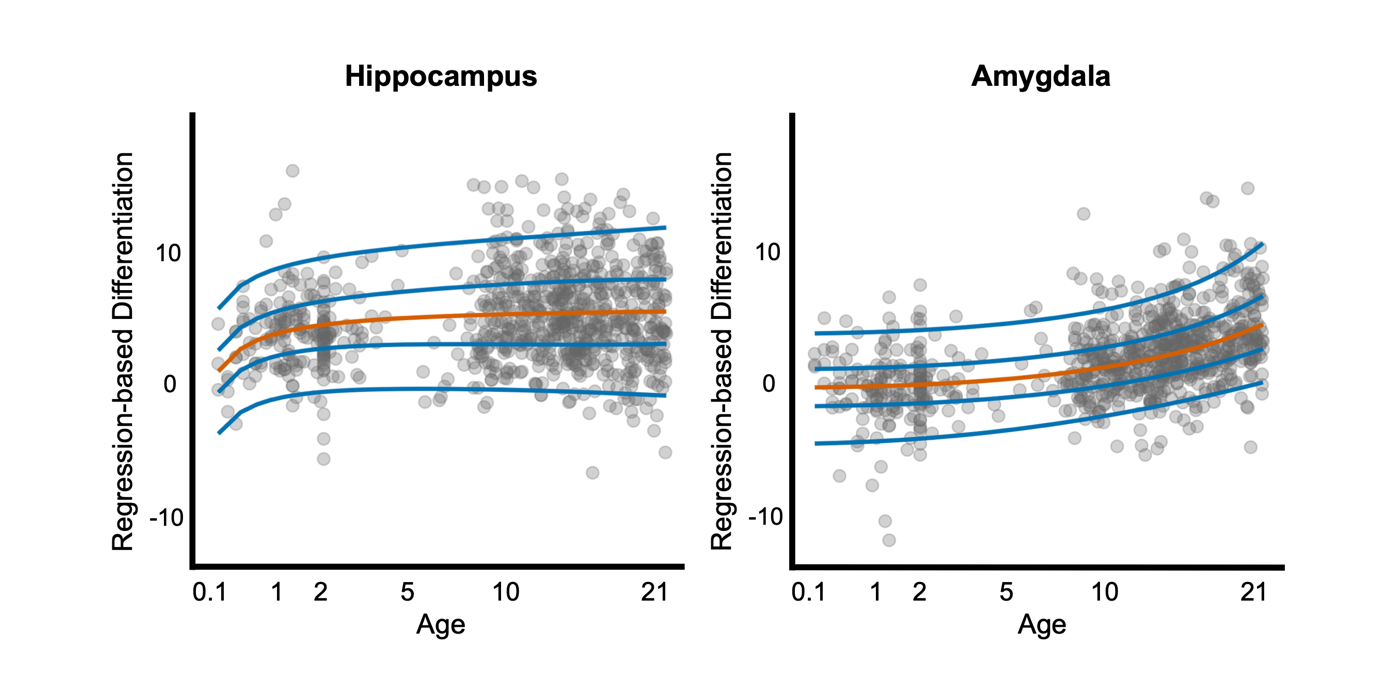

Figure S13. Normative Maturational Trajectory Using Regression-Based Differentiation Index. The age-related trajectories of differentiation are highly similar to the main results when using a differentiation index that takes into account whether at each voxel the connectivity is significantly differentiated.

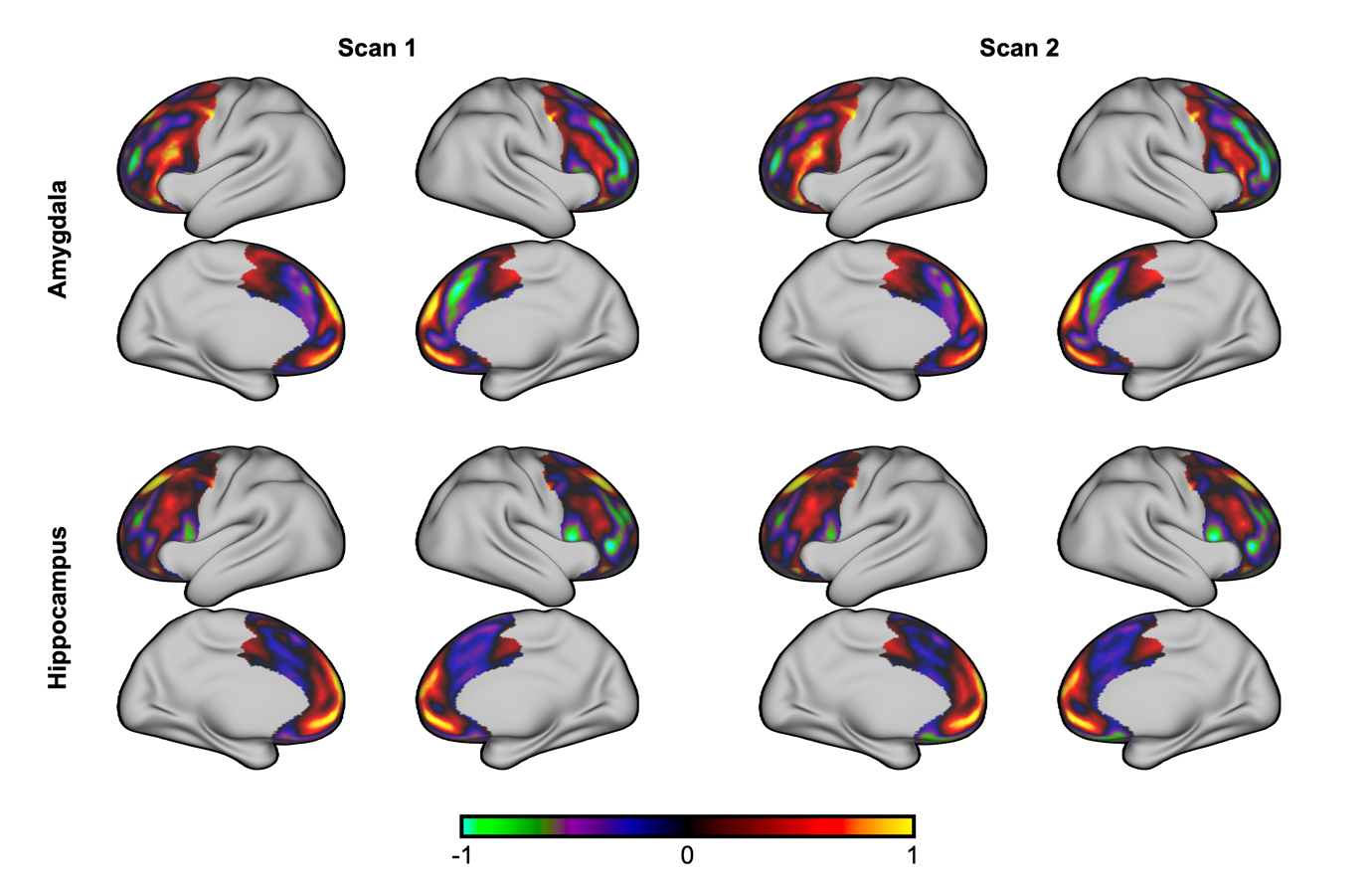

Figure S14. Reference Maps from Scan 1 and Scan 2. Across scans that took place at different days, group-average normalized connectivity patterns were highly similar for both fronto-amygdala and fronto-hippocampus connectivity.

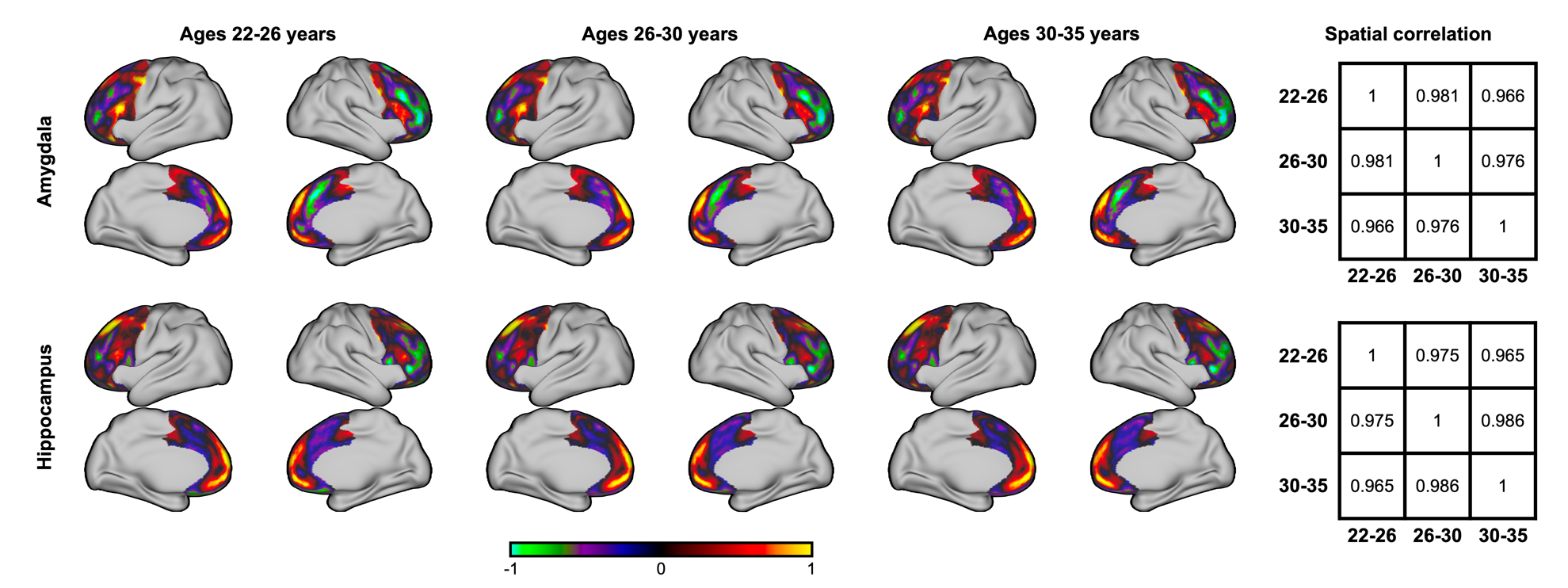

Figure S15. Reference Maps from Three Age-bins Across Young Adulthood. Across the three age-bins of 22-26, 26-30, and 30-35 across the young adult dataset, three sets of fronto-hippocampus and fronto-amygdala reference connectivity patterns are shown. Spatial correlation matrices report Pearson’s correlation values across the three age-bin connectivity patterns, respectively for hippocampus and amygdala.

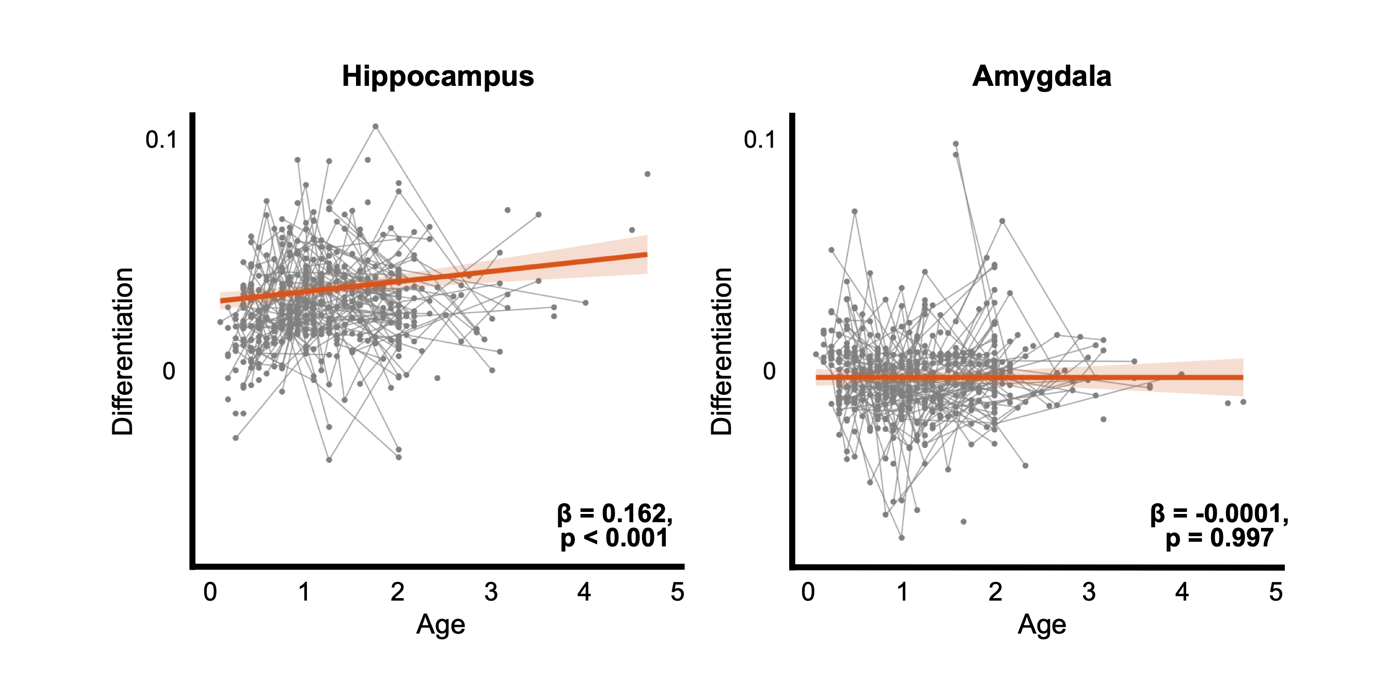

Figure S16. Age Trajectory of Frontolimbic Differentiation in BCP without Cross-Cohort Harmonization. Using all available scans from the BCP dataset (412 scans total), differentiation trajectories recapitulate what is seen in the main results where fronto-hippocampus connectivity shows early growth unlike the fronto-amygdala that is not developing in the early years of life. Linear mixed modeling was employed to account for longitudinal follow-up scans.

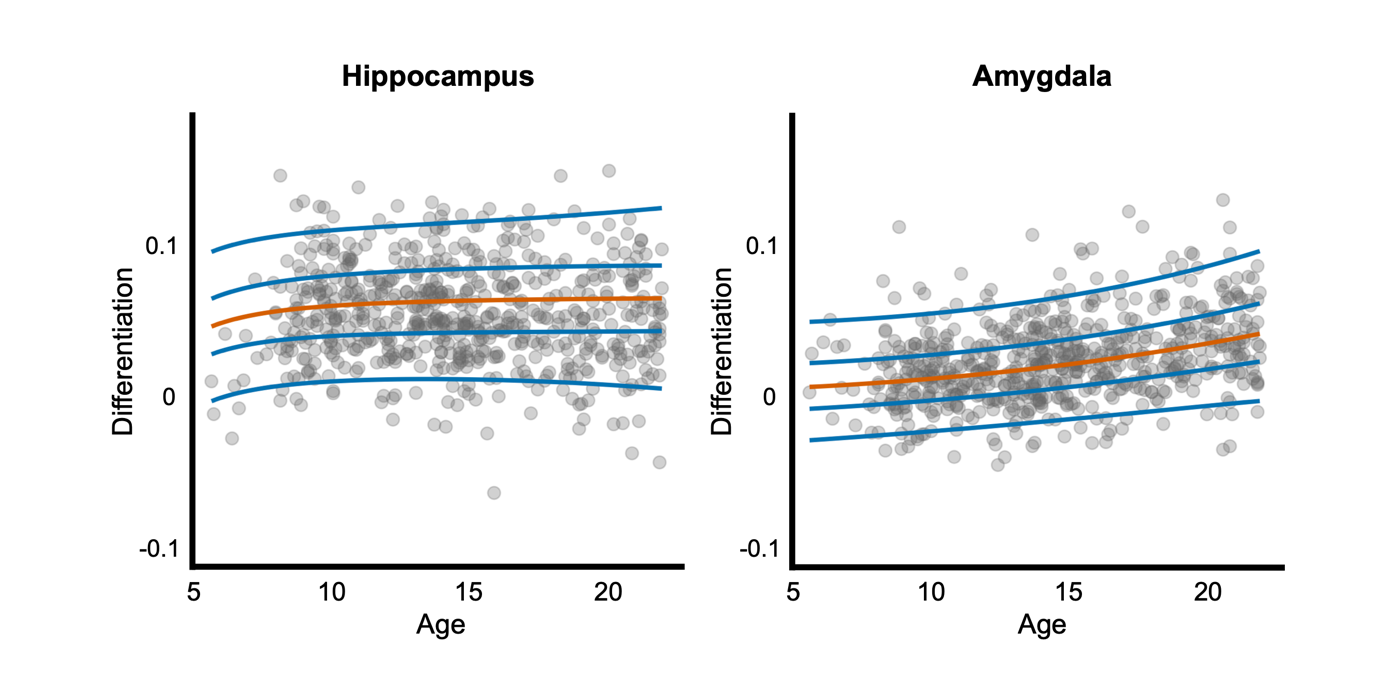

Figure S17. Age Trajectory of Frontolimbic Differentiation in HCP-D without Cross-Cohort Harmonization. Fitting the GAMLSS models with only the HCP-D dataset converges with the main results, with fronto-hippocampus differentiation plateauing across adolescence and fronto-amygdala differentiation maturing throughout adolescence into emerging adulthood. The same GAMLSS model distributions and parameters from the main analyses were applied here.

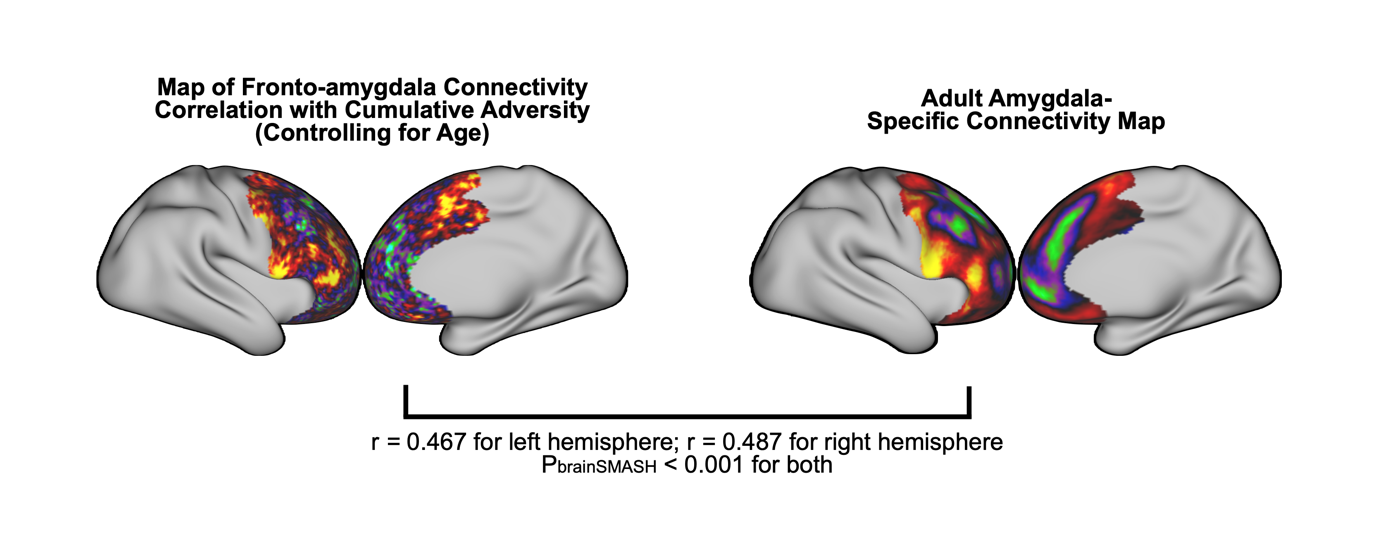

Figure S18. Correlation Pattern of Cumulative Adversity on Fronto-amygdala Connectivity. When controlling for age, fronto-amygdala connectivity correlation with cumulative adversity resembled the adult amygdala-specific connectivity map. In other words, as differentiation entails fronto-amygdala connectivity increases for amygdala-specific regions (positive, red in right map) and decreases for hippocampus-specific regions (negative, green in right map) as elaborated in the main results, cumulative adversity is associated with stronger increases and stronger decreases of the same regions at a given age. This indicates that cumulative adversity is associated with higher differentiation at a given age, which converges with the main results while demonstrating that this is an effect that exists across the frontal cortex.

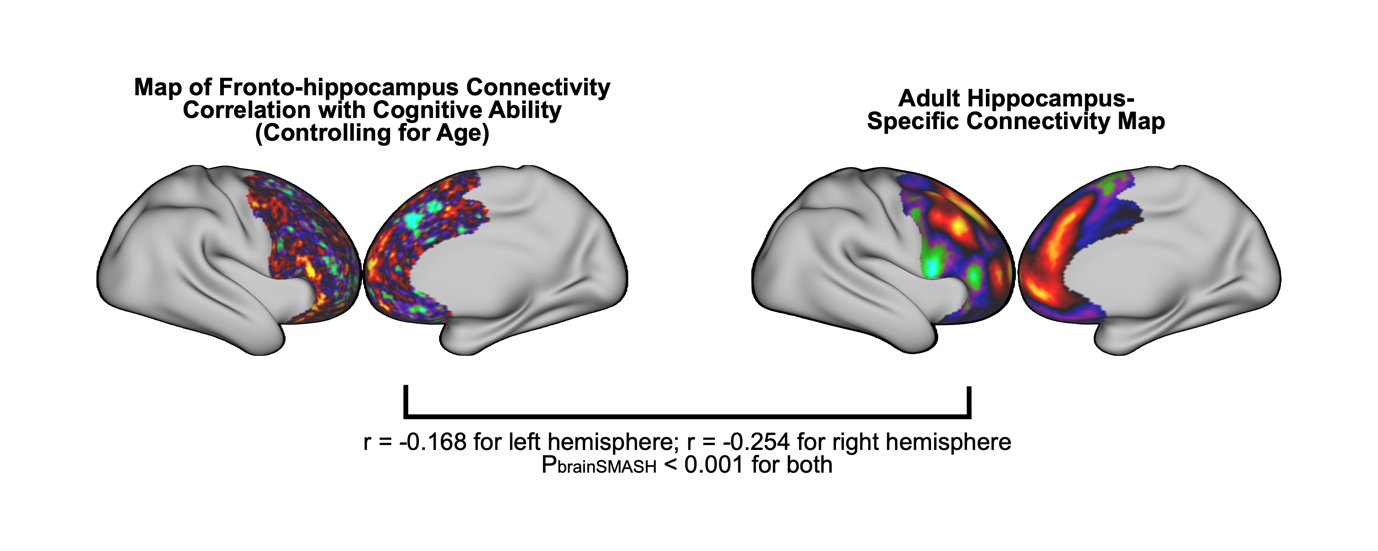

Figure S19. Correlation Pattern of Cognitive Ability on Fronto-hippocampus Connectivity. When controlling for age, fronto-hippocampus connectivity correlation with cognitive ability was anti-correlated with the adult hippocampus-specific connectivity map. Similar to Supplementary Figure 18, this implies that cognitive ability is associated with lesser differentiation of fronto-hippocampus connectivity across the frontal cortex, in line with the observation from the main results.

**Tables**

Table S1. Regions of Interest Definition. Cortical regions, hippocampus, and amygdala labels used to define the main regions of interest are listed. Region labels and the indexes are from the Glasser Atlas for cortical regions and the Melbourne Subcortex Atlas for hippocampus and amygdala.

| Cortical Region | No. | R_p10p_ROI | 186 | L_p10p_ROI | 366 | L_IFJp_ROI | 276 | R_10v_ROI | 104 | L_8BL_ROI | 266 |
| --- | --- | --- | --- | --- | --- | --- | --- | --- | --- | --- | --- |
| R_FEF_ROI | 26 | R_p47r_ROI | 187 | L_p47r_ROI | 367 | L_p9-46v_ROI | 279 | R_10pp_ROI | 106 | L_10d_ROI | 268 |
| R_PEF_ROI | 27 | L_FEF_ROI | 206 | R_47m_ROI | 82 | L_13l_ROI | 288 | R_25_ROI | 180 | L_10v_ROI | 284 |
| R_55b_ROI | 28 | L_PEF_ROI | 207 | R_8Av_ROI | 83 | L_47s_ROI | 290 | R_s32_ROI | 181 | L_10pp_ROI | 286 |
| R_6ma_ROI | 60 | L_55b_ROI | 208 | R_8C_ROI | 89 | L_s6-8_ROI | 294 | R_pOFC_ROI | 182 | L_s32_ROI | 361 |
| R_8Ad_ROI | 84 | L_6ma_ROI | 240 | R_44_ROI | 90 | R_SFL_ROI | 42 | R_a32pr_ROI | 195 | L_a32pr_ROI | 375 |
| R_9p_ROI | 87 | L_8Ad_ROI | 264 | R_45_ROI | 91 | R_24dv_ROI | 57 | R_p24_ROI | 196 | L_p24_ROI | 376 |
| R_a47r_ROI | 93 | L_9p_ROI | 267 | R_47l_ROI | 92 | R_SCEF_ROI | 59 | L_SFL_ROI | 222 | R_OFC_ROI | 109 |
| R_6r_ROI | 94 | L_a47r_ROI | 273 | R_IFJa_ROI | 95 | R_p24pr_ROI | 73 | L_24dv_ROI | 237 | L_OFC_ROI | 289 |
| R_IFJp_ROI | 96 | L_6r_ROI | 274 | R_p9-46v_ROI | 99 | R_33pr_ROI | 74 | L_SCEF_ROI | 239 | L_25_ROI | 360 |
| R_IFSp_ROI | 97 | L_IFSp_ROI | 277 | R_13l_ROI | 108 | R_a24pr_ROI | 75 | L_p24pr_ROI | 253 | L_pOFC_ROI | 362 |
| R_IFSa_ROI | 98 | L_IFSa_ROI | 278 | R_47s_ROI | 110 | R_p32pr_ROI | 76 | L_33pr_ROI | 254 | L_8BL_ROI | 266 |
| R_46_ROI | 100 | L_46_ROI | 280 | R_s6-8_ROI | 114 | R_a24_ROI | 77 | L_a24pr_ROI | 255 | Hippocampus and Amygdala | No. |
| R_a9-46v_ROI | 101 | L_a9-46v_ROI | 281 | L_47m_ROI | 262 | R_d32_ROI | 78 | L_p32pr_ROI | 256 | HIP-rh | 1 |
| R_9-46d_ROI | 102 | L_9-46d_ROI | 282 | L_8Av_ROI | 263 | R_8BM_ROI | 79 | L_a24_ROI | 257 | AMY-rh | 2 |
| R_9a_ROI | 103 | L_9a_ROI | 283 | L_8C_ROI | 269 | R_p32_ROI | 80 | L_d32_ROI | 258 | HIP-lh | 9 |
| R_a10p_ROI | 105 | L_a10p_ROI | 285 | L_44_ROI | 270 | R_10r_ROI | 81 | L_8BM_ROI | 259 | AMY-lh | 10 |
| R_11l_ROI | 107 | L_11l_ROI | 287 | L_45_ROI | 271 | R_9m_ROI | 85 | L_p32_ROI | 260 |  |  |
| R_6a_ROI | 112 | L_6a_ROI | 292 | L_47l_ROI | 272 | R_8BL_ROI | 86 | L_10r_ROI | 261 |  |  |
| R_i6-8_ROI | 113 | L_i6-8_ROI | 293 | L_IFJa_ROI | 275 | R_10d_ROI | 88 | L_9m_ROI | 265 |  |  |

Table S2. GAMLSS Distribution Choice. For both fronto-hippocampus and fronto-amygdala differentiation, all possible distributions were fitted before calculating the Akaike Information Criterion (AIC). Lower AIC indicates better fit, and the distributions with the lowest AIC were therefore used unless the model failed to converge in any cross-validation folds, model fit diagnostics were not satisfactory, or it was visually clear that the fitted model did not fit the data.

| Fronto-hippocampus Differentiation | | Fronto-amygdala Differentiation | |
| --- | --- | --- | --- |
| Distribution | AIC | Distribution | AIC |
| PE | -3371.0877 | ST2 | -3749.8174 |
| TF2 | -3369.4883 | SN1 | -3749.545 |
| TF | -3367.668 | ST1 | -3749.0301 |
| JSUo | -3366.8672 | JSUo | -3748.3427 |
| JSU | -3366.0339 | ST5 | -3746.4161 |
| ST3 | -3366.0164 | SEP4 | -3745.2216 |
| PE2 | -3365.7132 | EGB2 | -3745.2215 |
| ST2 | -3365.1957 | ST4 | -3744.8554 |
| SN2 | -3364.44 | SEP2 | -3744.7713 |
| ST1 | -3364.1206 | SHASH | -3744.5511 |
| SHASH | -3363.6248 | SST | -3744.1363 |
| ST5 | -3363.2939 | SHASHo | -3744.1098 |
| SHASHo2 | -3363.1581 | SHASHo2 | -3744.0981 |
| SN1 | -3362.7695 | ST3 | -3743.8639 |
| SHASHo | -3362.4215 | TF2 | -3743.5209 |
| SST | -3361.2618 | SEP3 | -3743.4285 |
| GT | -3359.8864 | TF | -3743.3752 |
| ST4 | -3355.3894 | PE | -3742.7695 |
| NET | -3328.7211 | SN2 | -3741.9497 |
|  |  | PE2 | -3741.3432 |
|  |  | GT | -3733.2985 |
|  |  | NET | -3715.9451 |

Table S3. Threshold-Free Cluster Enhancement (TFCE) Results in BCP. Six clusters in the hippocampus were found to be significant in their age-differentiation correlation. Clusters with very low number of voxels indicate that the effect was large enough to justify counting them as clusters as calculated by the TFCE algorithm.

| Cluster Index | Voxels | MAX | MAX X (vox) | MAX Y (vox) | MAX Z (vox) | COG X (vox) | COG Y (vox) | COG Z (vox) |
| --- | --- | --- | --- | --- | --- | --- | --- | --- |
| 6 | 8 | 0.992 | 32 | 55 | 28 | 32.7 | 54.5 | 28.5 |
| 5 | 7 | 0.998 | 34 | 55 | 25 | 33.6 | 55 | 25.7 |
| 4 | 5 | 0.992 | 56 | 52 | 28 | 56.6 | 53.2 | 28.2 |
| 3 | 2 | 0.993 | 61 | 50 | 30 | 61 | 49.5 | 30 |
| 2 | 2 | 0.972 | 59 | 47 | 31 | 59 | 47.5 | 31 |
| 1 | 1 | 0.99 | 58 | 44 | 35 | 58 | 44 | 35 |

Table S4. Threshold-Free Cluster Enhancement Results in HCP-D. Three clusters of voxels in the amygdala were significant in their age-differentiation correlation.

| Cluster Index | Voxels | MAX | MAX X (vox) | MAX Y (vox) | MAX Z (vox) | COG X (vox) | COG Y (vox) | COG Z (vox) |
| --- | --- | --- | --- | --- | --- | --- | --- | --- |
| 3 | 171 | 1 | 59 | 59 | 25 | 56.7 | 60.3 | 25.7 |
| 2 | 120 | 1 | 34 | 63 | 27 | 32.7 | 61.1 | 26.1 |
| 1 | 1 | 0.959 | 31 | 52 | 28 | 31 | 52 | 28 |

Table S5. Spatial Permutation Test of BCP Subregion Significance. Spatially permuting the t-statistic value maps of the age-related differentiation effect yielded p-values for whether the observed effect is localized to certain subregions. Lack of significance across all subregions indicate the effect was not localized to any hippocampus subregion in the BCP.

| Subregion | Observed Sum of T-statistic value | Permutation p-value |
| --- | --- | --- |
| Right-Anterior Hippocampus | 293.07 | 0.0898 |
| Right-Posterior Hippocampus | 66.12 | 0.3099 |
| Left-Anterior Hippocampus | 225.90 | 0.1074 |
| Left-Posterior Hippocampus | 285.84 | 0.0832 |

Table S6. Spatial Permutation Test of HCP-D Subregion Significance. Lack of significance across all subregions indicate the effect was not localized to any amygdala subregion in the HCP-D.

| Subregion | Observed Sum of T-statistic value | Permutation P-value |
| --- | --- | --- |
| Right-Medial Amygdala | 213.16 | 0.0842 |
| Right-Lateral Amygdala | 325.41 | 0.0768 |
| Left-Medial Amygdala | 230.04 | 0.0856 |
| Left-Lateral Amygdala | 270.39 | 0.0834 |

Table S7. Multiple Regression Model Statistics for the Spatial Axes. X, Y, and Z axes each indicate the medial-lateral, posterior-anterior, and dorsal-ventral axes. Absolute values were used for the X axis to represent medial-to-lateral direction for both left and right hemispheres.

| **Region** | **Term** | **Estimate** | **Standard Error** | **t-statistic** | **p-value** |
| --- | --- | --- | --- | --- | --- |
| Hippocampus | Intercept | 0.0010 | 0.0002 | 6.1550 | <0.0001 |
|  | abs(X) | 0.0004 | 0.0002 | 2.3529 | 0.0188 |
|  | Y | 0.0020 | 0.0004 | 4.6345 | <0.0001 |
|  | Z | 0.0006 | 0.0004 | 1.4830 | 0.1383 |
| Amygdala | Intercept | 0.0069 | 0.0002 | 28.4924 | <0.0001 |
|  | abs(X) | 0.0007 | 0.0003 | 2.9588 | 0.0032 |
|  | Y | 0.0011 | 0.0003 | 3.6439 | 0.0003 |
|  | Z | 0.0006 | 0.0003 | 1.9059 | 0.0572 |

Table S8. Item Importance of Adversity Association. Feature weights from the Principal Component Regression model signify the importance of individual adversity items. Bootstrapping was used to estimate the 95% confidence intervals (CI), and p-values were derived from the bootstrapping results. Statistical significance was assessed using p-values adjusted to control the False Discovery Rate (FDR).

| Feature | Importance | CI Lower | CI Upper | p-value | FDR-corrected p-value |
| --- | --- | --- | --- | --- | --- |
| Parent went to jail | 0.0417 | 0.0033 | 0.1025 | 0.0058 | 0.1450 |
| Fam mem arrest | 0.0291 | -0.0045 | 0.0629 | 0.0712 | 0.4450 |
| Attended new school | 0.0281 | -0.0036 | 0.1053 | 0.1176 | 0.5129 |
| Mother father lost job | 0.0280 | -0.0004 | 0.0682 | 0.0546 | 0.4450 |
| Parent new job | 0.0242 | -0.0008 | 0.0822 | 0.0602 | 0.4450 |
| Parents financial situation | 0.0221 | -0.0188 | 0.0570 | 0.1738 | 0.5431 |
| Victim crime assault | 0.0218 | -0.0081 | 0.0730 | 0.1436 | 0.5129 |
| Parent away home | 0.0199 | -0.0065 | 0.0723 | 0.1414 | 0.5129 |
| Parents trouble law | 0.0158 | -0.0147 | 0.0505 | 0.2532 | 0.6086 |
| Family moved | 0.0144 | -0.0151 | 0.0466 | 0.2460 | 0.6086 |
| Fam mem injured | 0.0121 | -0.0183 | 0.0461 | 0.2678 | 0.6086 |
| Fam mem died | 0.0111 | -0.0179 | 0.0556 | 0.3200 | 0.6667 |
| Fam mem drug alc | 0.0089 | -0.0298 | 0.0376 | 0.4008 | 0.7708 |
| Parents sep 12 months | -0.0074 | -0.0641 | 0.0250 | 0.8822 | 0.9818 |
| Saw crime accident | 0.0065 | -0.0292 | 0.0368 | 0.4924 | 0.8150 |
| Lost close friend | 0.0064 | -0.0244 | 0.0509 | 0.6194 | 0.8150 |
| New stepparent | 0.0059 | -0.0270 | 0.0327 | 0.6462 | 0.8150 |
| Fam mem emotional mental | 0.0052 | -0.0296 | 0.0483 | 0.6250 | 0.8150 |
| Close friend died | 0.0040 | -0.0299 | 0.0317 | 0.5906 | 0.8150 |
| Sick injured | 0.0034 | -0.0401 | 0.0312 | 0.6520 | 0.8150 |
| Sibling left home | 0.0026 | -0.0427 | 0.0304 | 0.6502 | 0.8150 |
| Parents divorced | -0.0011 | -0.0428 | 0.0297 | 0.9818 | 0.9818 |
| Parents argue | -0.0011 | -0.0365 | 0.0388 | 0.9778 | 0.9818 |
| Close friend sick injured | -0.0005 | -0.0547 | 0.0306 | 0.7760 | 0.9238 |
| New sibling | 0.0002 | -0.0326 | 0.0350 | 0.9504 | 0.9818 |

Table S9. Domain Importance of Cognition Association. Feature weights from the Principal Component Regression model signify the importance of individual cognitive domains. Bootstrapping was used to estimate the 95% confidence intervals (CI), and p-values were derived from the bootstrapping results. Statistical significance was assessed using p-values adjusted to control the False Discovery Rate (FDR).

| Feature | Importance | CI Lower | CI Upper | p-value | FDR-corrected p-value |
| --- | --- | --- | --- | --- | --- |
| Picture Vocabulary | 0.0479 | 0.0147 | 0.0877 | 0.0056 | 0.0336 |
| List Sorting Working Memory | 0.0450 | 0.0112 | 0.1077 | 0.0096 | 0.0336 |
| Oral Reading Recognition | 0.0439 | 0.0091 | 0.0824 | 0.0184 | 0.0429 |
| Pattern Comparison Processing Speed | 0.0405 | -0.0029 | 0.1272 | 0.072 | 0.126 |
| Dimensional Change Card Sort | 0.0279 | -0.0196 | 0.0680 | 0.2268 | 0.2646 |
| Picture Sequence Memory | 0.0240 | -0.0212 | 0.0678 | 0.1308 | 0.1831 |
| Flanker Inhibitory Control and Attention | 0.0105 | -0.0719 | 0.0549 | 0.5784 | 0.5784 |

Table S10. Behavior Measures in the PNC. Cumulative trauma was calculated by summing all scores across the K-SADS PTSD module items. Cognitive ability was calculated by z-scoring and summing test scores across diverse cognitive domains.

Cumulative Trauma: K-SADS PTSD module

Have you ever been in a flood or a tornado or an earthquake or a hurricane or some other natural disaster where you thought you were going to die or be seriously hurt?

Have you ever been in a situation where you thought you or someone close to you was going to be killed or be hurt very badly (e.g. family violence)?

Have you ever been attacked by somebody or badly beaten?

Have you ever been very upset by someone forcing you to do something sexual?

Have you ever been attacked sexually or raped?

Have you ever been threatened with a weapon?

Have you ever been in a bad accident?

Other than television or at the movies, have you ever seen or heard somebody get killed or get hurt very badly or die?

Have you ever been very upset by seeing a dead body or by seeing pictures of the dead body of somebody you knew well?

Cognitive Ability: Cognitive Test Scores

Penn Continuous Performance Test – Number of Correct Responses to Number Trials (TP), Number of Incorrect Responses to Number Trials (FP - Inverted)

Penn Verbal Reasoning Test – Total Correct Responses for All Test Trials, Median Response Time for Correct Verbal Reasoning Responses (Inverted)

Wide Range Assessment Test Total Standard Score

Penn Line Orientation Test – Percent Correct Responses for All Test Trials, Median Response Time for Correct Trials (Inverted)

Penn Matrix Analysis Test – Percent of Correct Responses for All Test Trials, Median Response Time for Correct Test Trial Responses (Inverted)

Penn Conditional Exclusion Test – Calculated Accuracy Measure, Median Response Time for Correct Responses

Penn Face Memory Test – Number of Correct Responses to Target Faces (TP), Number of Incorrect Responses to Foil Faces (FP - Inverted)

Penn Word Memory Test – Number of Correct Responses to Target Words (TP), Number of Incorrect Responses to Foil Words (FP)

Penn Visual Object Memory Test – Number of Correct Responses to Target Shapes (TP), Number of Incorrect Responses to Foil Shapes (FP - Inverted)

Letter N-Back – Number of Correct Responses to for 1-Back and 2-Back Trials, Mean of the Median Response Time for Correct Responses for 1-Back (TP) and for 2-Back (TP) Trials (Inverted)

Penn Emotion Identification Test – Total Correct Responses for All Test Trials, Median Response Time for Total Correct Test Trial Responses (Inverted)

Table S11. Reference Map Similarity in 100 Random Half-Splits. Repeated half-splits within the HCP-YA dataset scan 1 resulted in a distribution of within-dataset reference map correlations. The high correlation values imply that the reference maps are good approximations of population-level mean, not driven by certain groups of individuals specific to the HCP-YA dataset.

| Frontal Cortex Connectivity Map | Half-Split Correlation Mean | Half-Split Correlation Standard Deviation |
| --- | --- | --- |
| Left Hippocampus | 0.9858 | 0.0012 |
| Left Amygdala | 0.9644 | 0.0015 |
| Right Hippocampus | 0.9872 | 0.0013 |
| Right Amygdala | 0.9779 | 0.0013 |

**Supporting Movies**

Movie S1. Hippocampus Connectivity Pattern Across Medial Frontal Cortex. Across maturation, spatial reorganization of fronto-hippocampus connectivity is shown for medial frontal cortex.

Movie S2. Amygdala Connectivity Pattern Across Medial Frontal Cortex. Across maturation, spatial reorganization of fronto-amygdala connectivity is shown for medial frontal cortex.

Movie S3. Hippocampus Connectivity Pattern Across Lateral Frontal Cortex. Across maturation, spatial reorganization of fronto-hippocampus connectivity is shown for lateral frontal cortex.

Movie S4. Amygdala Connectivity Pattern Across Lateral Frontal Cortex. Across maturation, spatial reorganization of fronto-amygdala connectivity is shown for lateral frontal cortex.
